# Predicting brain-age from multimodal imaging data captures cognitive impairment

**DOI:** 10.1101/085506

**Authors:** Franziskus Liem, Gaël Varoquaux, Jana Kynast, Frauke Beyer, Shahrzad Kharabian Masouleh, Julia M. Huntenburg, Leonie Lampe, Mehdi Rahim, Alexandre Abraham, R. Cameron Craddock, Steffi Riedel-Heller, Tobias Luck, Markus Loeffler, Matthias L. Schroeter, Anja Veronica Witte, Arno Villringer, Daniel S. Margulies

## Abstract

The disparity between the chronological age of an individual and their brain-age measured based on biological information has the potential to offer clinically-relevant biomarkers of neurological syndromes that emerge late in the lifespan. While prior brain-age prediction studies have relied exclusively on either structural or functional brain data, here we investigate how multimodal brainimaging data improves age prediction. Using cortical anatomy and whole-brain functional connectivity on a large adult lifespan sample (N = 2354, age 19-82), we found that multimodal data improves brain-based age prediction, resulting in a mean absolute prediction error of 4.29 years. Furthermore, we found that the discrepancy between predicted age and chronological age captures cognitive impairment. Importantly, the brain-age measure was robust to confounding effects: head motion did not drive brain-based age prediction and our models generalized reasonably to an independent dataset acquired at a different site (N = 475). Generalization performance was increased by training models on a larger and more heterogeneous dataset. The robustness of multimodal brain-age prediction to confounds, generalizability across sites, and sensitivity to clinically-relevant impairments, suggests promising future application to the early prediction of neurocognitive disorders.

**Highlights:**

- Brain-based age prediction is improved with multimodal neuroimaging data.
- Participants with cognitive impairment show increased brain aging.
- Age prediction models are robust to motion and generalize to independent datasets from other sites.

## 1. Introduction

The brain continues to change throughout adult life. Structural aspects, such as cortical thinning, demonstrate robust patterns of alteration during adulthood (Hogstrom et al., 2013; Storsve et al., 2014). Likewise, age-related differences in brain function, demonstrated through studies of functional connectivity, have also been observed (Damoiseaux et al., 2008; Dennis & Thompson, 2014).

Establishing the trajectories of such changes over the lifespan provides a basis for characterizing clinically-relevant deviations (Ziegler et al., 2012; Raz & Rodrigue, 2006). Brain-based age prediction offers a promising approach for providing personalized biomarkers of future cognitive impairments by capturing deviations from typical development of brain structure and function.

Brain-based age prediction aims to estimate a person’s age based on brain data acquired using magnetic resonance imaging (MRI, Franke et al., 2010; Franke & Gaser, 2012). In a first step, an age prediction model is trained based on brain imaging data from a large lifespan sample. In a second step, this model can be used to estimate a novel individual’s age based solely on their brain-imaging data. By comparing a person’s estimated age with their chronological age, conclusions about age-typical and atypical brain development can be drawn.

Brain-based age prediction exemplifies a larger trend in neuroscience (Bzdok, 2016; Gabrieli et al., 2015; Pereira et al., 2009; Varoquaux & Thirion, 2014) and psychology (Yarkoni & Westfall, 2016) to move from correlative to predictive studies, often using tools from machine learning. Individual brain-based prediction and classification may give rise to brain imaging-based biomarkers that could aid clinical diagnostics, for instance, by predicting an individual’s risk of developing dementia based on their brain (Bron et al., 2015).

One successful age prediction framework is based on structural brain data analyzed with voxel-based morphometry (VBM, Franke et al., 2010; Franke & Gaser, 2012). Using this approach, accelerated brain aging was found in patients with Alzheimer’s disease (Franke et al., 2010; Franke & Gaser, 2012), traumatic brain injuries (Cole et al., 2015), psychiatric disorders (Koutsouleris et al., 2013), and subjects with risks to physical health (Franke et al., 2014). This brain-age metric can also predict the future conversion from mild cognitive impairment to Alzheimer’s disease (Gaser et al., 2013). This computational approach is not restricted to showing accelerated brain aging as a negative effect but has also been used to demonstrate the positive effects of education, physical exercise (Steffener et al., 2016), and meditation (Luders et al., 2016) on brain aging. Other work has shown that accelerated brain development is related to accelerated cognitive development in young subjects (Erus et al., 2014).

In addition to brain structure, functional connectivity based on resting-state fMRI data (Craddock et al., 2013) also has the potential to provide clinically-relevant biomarkers (Craddock et al., 2009; Castellanos et al., 2013), as the data is easily acquired in a clinical setting (Greicius, 2008; Damoiseaux et al., 2012). Similar to the structural age estimation approach, Dosenbach et al. (2010) demonstrated that this is also feasible with resting-state functional connectivity data from young subjects. As different MRI modalities capture not only shared but also unique information about brain aging (Groves et al., 2012), prediction accuracy may benefit by incorporating these additional sources of information. For instance, Brown et al. (2012) and Erus et al. (2014) have shown that combining information from gray and white matter anatomy increases prediction accuracy in young subjects. The present study investigates how combining data from two even more dissimilar sources, brain anatomy and functional connectivity, influences age prediction in a lifespan sample. This is important as function and structure convey converging as well as diverging information (Damoiseaux & Greicius, 2009).

While machine-learning methods enable predictions on a single-subject level, factors driving these predictions are often difficult to determine. Predictions that appear to be based on brain information may actually be driven by confounds. One major confound in functional and structural MRI is head motion (Satterthwaite et al., 2013; Power et al., 2012; Reuter et al., 2015; Alexander-Bloch et al., 2016). For instance, head motion can make cortex appear thinner (Reuter et al., 2015). An age-related increase in head motion might give rise to a supposedly ‘brain-based’ age predictor that relies heavily on head motion. Furthermore, while machine learning models are trained on one dataset and evaluated on another, in neuroimaging these datasets often come from the same study, i.e., same site and scanner. In such cases, models may overfit one site’s subtle idiosyncrasies, rendering poor predictive power for data from another site. Therefore, in the current study we aimed to address these confounds by determining the effect of head motion on brain-based age prediction and predictive performance on data from a novel site.

The present study investigates (i) whether incorporating multiple imaging modalities increases prediction accuracy, (ii) whether cognitive impairments are related to brain aging, and (iii) how robust our predictive models are, specifically regarding head motion and generalizabilty to new datasets. Using data from brain anatomy and functional connectivity, we show that (i) incorporating multiple modalities increases predictive performance, (ii) cognitive impairments are related to advanced brain aging, and (iii) our models are robust as they are not driven by head motion and generalize reasonably to new datasets.

## 2. Materials: lifespan data & preprocessing

Two independent samples were investigated in this study: the LIFE (Loeffler et al., 2015) and the Enhanced Nathan Kline Institute – Rockland sample (NKI, Nooner et al., 2012). Since the majority of analyses is performed on the LIFE dataset, the NKI set is described in detail in Appendix A.

### 2.1. LIFE sample

Participants took part in the LIFE-Adult-Study (life.uni-leipzig.de, Loeffler et al., 2015) of the Leipzig Research Centre for Civilization Diseases (LIFE) and were randomly selected, community-dwelling volunteers. The study was approved by the institutional ethics board of the Medical Faculty of the University of Leipzig. Participants signed an informed consent form and were paid for their participation.

Of the 10000 volunteers participating in the LIFE-study, approximately 2600 also underwent MRI assessment and neuropsychological testing. In the present study, data from 2354 individuals were included. Exclusion criteria included neuroradiological findings, missing MRI data or excessive head motion in the functional scans (mean FD > 0.6 mm, Power et al., 2012). Subjects were between 19 and 82 years (*M* = 58.68; *S D* = 15.17; 1120 female, 1234 male).

### 2.2. Cognitive phenotyping

#### Neurocognitive Disorder

The Diagnostic Statistical Manual of Mental Disorders 5th edition (DSM-5, American Psychiatric Association, 2013) introduced Neurocognitive Disorder (NCD) as a new diagnostic category for acquired cognitive dysfunction. NCD diagnosis comprises the evaluation of subjective cognitive complaints, cognitive performance and independence in activities of daily living. To this end, a comprehensive and domain specific neuropsychological evaluation is required, including the cognitive domains attention, executive function, memory, language, visuoconstruction, and social cognition.

Two subtypes of NCD are distinguished: mild and major NCD. Both sub-types are characterized by subjective cognitive complaints. Mild NCD is presented with a cognitive performance decline that ranges between -1 and -2 SD below age and gender norms in at least one cognitive domain, and preserved independence in daily life. Contrary, persons with major NCD have severe cognitive deficits (<-2 SD below age and gender norms) in at least one cognitive domain that interfere with independence in everyday activities. Thus, the term major NCD represents the current concept of dementia.

The DSM-5 criteria for NCD diagnosis served as a template to characterize the study sample with respect to objective cognitive impairment (OCI). Only OCI was investigated in the present study. Independence in daily activities and subjective cognitive complaints were not considered as a criterion for cognitive phenotyping in this study as consensus questionnaires are still lacking.

#### Neuropsychological assessment

Cognitive performance was assessed with a set of standard neuropsychological tests, spanning several cognitive domains (Loeffler et al., 2015). All scores were carefully checked for missing values and plausibility. The Stroop test (Stroop, 1935; Treisman & Fearnley, 1969; Zysset et al., 2001; Schroeter et al., 2002), which quantifies executive functions, was administered. Social cognition was assessed with the Reading the Mind in the Eyes test (RMET, Cohen et al., 2001; Bölte, 2005), which quantifies the ability to infer mental states only from eye gaze. The CERAD-plus (Thalmann et al., 1997; Morris et al., 1989) is a dementia screening battery focusing on Alzheimer’s disease. It includes tests of verbal and figural memory and learning (10-items word list, figure recall), language (Boston Naming Test, semantic and phonematic verbal fluency), and visuoconstrution (figure copy). This battery also includes the Mini Mental State Examination (MMSE, Folstein et al., 1975) as well as the Trail Making Test (TMT, Reitan, 1979), which measures visual attention and cognitive flexibility.

#### Domain-specific scores

For the domain-specific evaluation of cognitive performance, test scores were assigned to the cognitive domains proposed by the DSM-5 and aggregated (Beck et al., 2014).

Scores for the following domains were calculated:

- *attention* (TMT-A time to complete (TTC), TMT-A errors, Stroop neutral reaction time (RT), Stroop neutral % correct),
- *executive function* (TMT-B TTC / TMT-A TTC, Stroop incongruent TTC / Stroop neutral TTC),
- *memory* (CERAD word list (trial 1, 2, 3, total, delayed recall, recognition), CERAD figure delayed recall),
- *language* (Boston Naming Test, semantic fluency (animals), phonematic fluency (s-words)),
- *visuoconstruction* (CERAD figure copy), and
- *social cognition* (RMET).

#### Objective cognitive impairment (OCI)

To translate neuropsychological measures into scores informative of cognitive performance independent of age, sub-test scores were z-standardized within the corresponding ageand sex group (18-39, 40-49, 50-59, 60-64, 65-69, 70-74, 75+ years). Where necessary, standardized scores were inverted so that a higher score reflects higher performance. If more than one sub-test measure per domain was available, sub-tests were averaged within domains. These domain-specific scores can be interpreted as the average performance of a person in a certain domain and a deviation from their peers’ performance in that domain. Based on the domain-specific scores and in analogy to aforementioned NCD classification scheme, subjects were classified as *OCI-norm*, - *mild* (at least one domain score between -1 and -2 SD), or - *major* (at least one domain score <-2 SD), if they had at least three valid domain scores.

### 2.3. MR data

Brain imaging was performed using a 3T Siemens Trio scanner equipped with a 32 channel head coil.

Resting-state functional images were acquired using an T2*-weighted echo-planar imaging sequence with an in-plane voxel size of 3×3 mm, slice thickness of 4 mm, slice gap of 0.8 mm, 30 slices, echo time (TE) of 30 ms, repetition time (TR) of 2000 ms and a flip-angle of 90°. This sequence lasted 10 min (300 volumes), during which participants were instructed to keep their eyes open and not to fall asleep. A gradient-echo fieldmap with the same geometry was recorded for distortion correction (TR = 488 ms, TE 1 = 5.19 ms, TE 2 = 7.65 ms).

High resolution T1-weighted structural images were acquired using an MP-RAGE sequence with 1 mm isotropic voxels, 176 slices, TR = 2300 ms, TE = 2.98 ms, and inversion time (TI) = 900 ms.

### 2.4. MR data preprocessing

MRI data processing was implemented in a python pipeline via Nipype (v0.10.0, Gorgolewski et al., 2011), which included routines from FSL (v5.0.9, Jenkinson et al., 2012), FreeSurfer (v5.3, Fischl, 2012), ANTS (v2.1.0, Avants et al., 2011), CPAC (`v0.3.9.1, fcp-indi.github.io`), and Nilearn (`v0.2.3, nilearn.github.io`, Abraham et al., 2014). The pipeline is available at github.com/fliem/LIFE_RS_preprocessing.

#### Functional MRI

After removal of the first five volumes (to allow the magnetization to reach a steady state) and motion correction (FSL mcflirt), rigid body coregistration of the functional scan to the anatomical image (FreeSurfer bbregister), as well as EPI distortion corrections (FSL fugue) were calculated and jointly applied in a subsequent step to each volume of the functional scan. Denoising included removal of (i) 24 motion parameters (CPAC, Friston et al., 1996), (ii) motion and signal intensity spikes (Nipype rapidart), (iii) six components explaining the highest variance from a singular value decomposition of white matter and cerebrospinal fluid time series (CompCor, Behzadi et al., 2007, signals extracted from individual masks created with FSL fast, decomposition executed with CPAC), and (iv) linear and quadratic signal trends. Subsequently, functional data were morphed to MNI space via transformation fields estimated from the structural data (ANTS). Functional data were then band-pass filtered between 0.01 and 0.1 Hz (Nilearn).

#### Structural MRI

The FreeSurfer software package was used to create models of the cortical surface for cortical thickness and cortical surface area measurements. Subcortical volumes were obtained from the automated procedure for volumetric measures of brain structures implemented in FreeSurfer.

## 3. Methods: age prediction analysis

### 3.1. Age prediction

Models were trained to predict age based on a variety of input data, i.e., functional connectomes of two different spatial resolutions and measures of cortical anatomy (cortical thickness, cortical surface area, subcortical volumes). A schematic overview of the age prediction analysis is shown in Figure 1.

**Figure 1.**
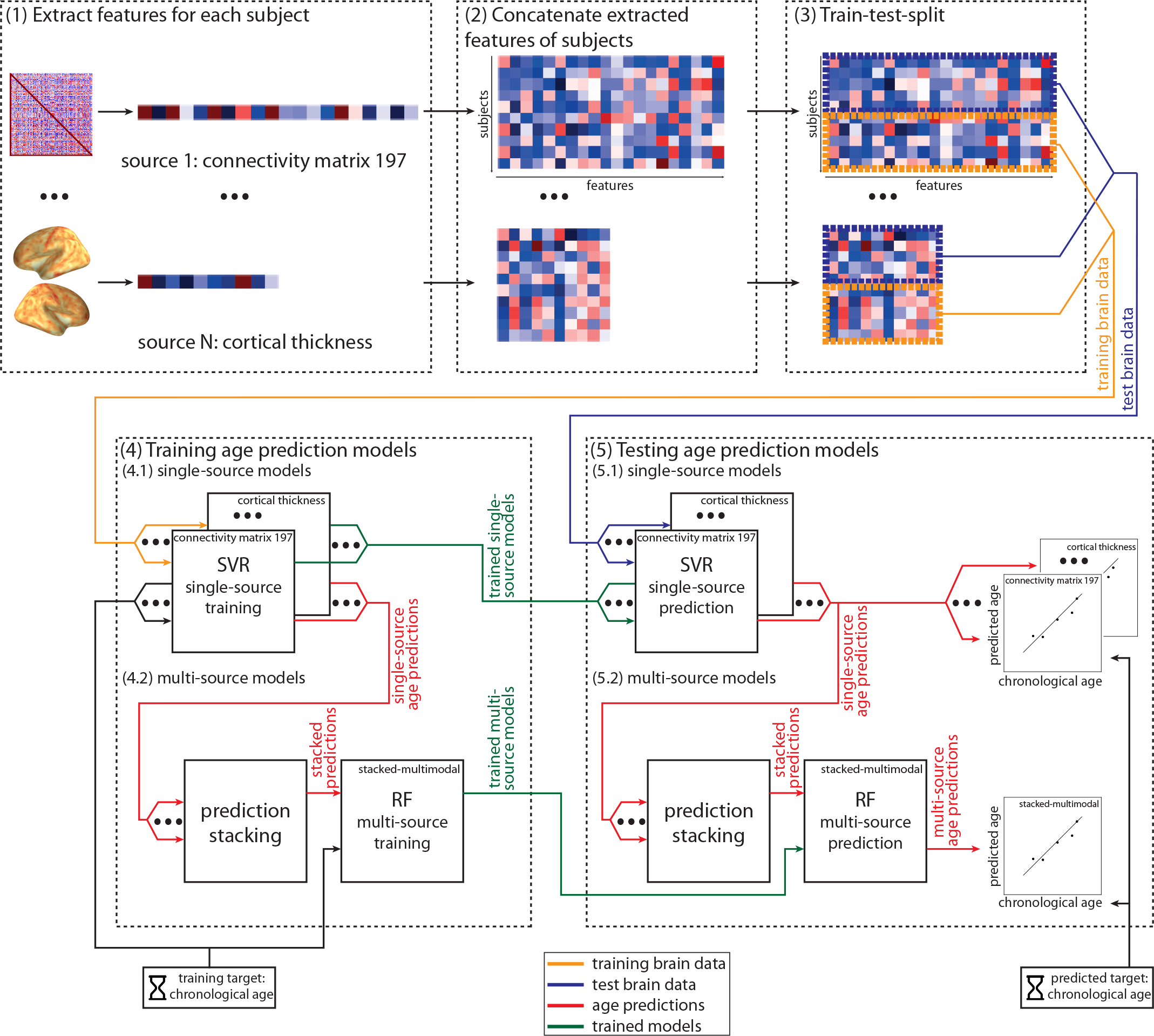
Overview of feature extraction and predictive analysis. (1) For each subject, feature vectors from the following data sources are extracted: *connectivity matrix 197, connectivity matrix 444, cortical thickness, cortical surface area,* and *subcortical volumes*. (2) Within each source, data for subjects are concatenated to obtain input data matrices. (3) Data is split into training and test set. (4) Training data (yellow line) is used to train age prediction models. (4.1) First, five singlesource support vector regression models (SVR) are trained to predict chronological age based on training brain data. (4.2) Second, the single-source predictions (red line) are stacked and entered into the training of multi-source random forest models (RF). Three separate multi-source models were trained: *stacked-function* (combining *connectivity matrix 197* and *connectivity matrix 444*), *stacked-anatomy* (combining *cortical thickness, cortical surface area, and subcortical volumes*), and *stacked-multimodal* (combining all five single-source models). (5) The trained models (green line) are then evaluated. (5.1) Trained single-source models give single-source age predictions based on test data (blue line). (5.2) These predictions are stacked and entered into the trained multi-source models to obtain multi-source age predictions. Prediction performance is evaluated by comparing predicted age with chronological age.

#### 3.1.1. Input data

The following five sources of neuroimaging data entered the age prediction models. Two sources represent brain connectivity in different spatial resolutions, three sources originate from brain anatomy:

1. *connectivity matrix 197,*
2. *connectivity matrix 444,*
3. *cortical thickness,*
4. *cortical surface area,* and
5. *subcortical volumes*

After extracting feature vectors for each subject and modality (see Figure 1.1), vectors were stacked to obtain the input data matrices for the age prediction analysis (see Figure 1.2).

*Brain function*. Functional connectomes were derived from preprocessed functional MRI data using the Nilearn package. Mean time-series were extracted from cortical and subcortical regions of the functionally defined BASC parcellation atlas (Bellec et al., 2010, obtained via the Nilearn data fetcher fetch_atlas_basc_multiscale_2015). Functional connectivity between all pairs of regions was quantified via Pearson correlation, resulting in a symmetric connectivity matrix. Since measures derived from connectomes vary with parcellation resolution (Fornito et al., 2010) and there is no ‘right’ number of parcels, we investigated two different levels of spatial granularity. Based on Thirion et al. (2014), who recommend parcellations consisting of around 200 to 500 regions, we reconstructed connectivity matrices from 197 and 444 regions.

Connectivity matrices underwent Fisher’s r-to-z transformation and a feature vector was extracted from the lower triangle (*N*_*features*_(*connectivity matrix 197*) = 19306, *N*_*features*_(*connectivity matrix 444*) = 98346). The shape of the input matrix was *N*_*subjects*_ × *N*_*features*_ (with *Nsubjects* varying between analyzes; see section 3.2).

*Brain anatomy*. Native surface models for cortical thickness and surface area were transformed into the fsaverage4 standard space. The data for the two hemispheres was concatenated (*N*_*features*_(*cortical thickness*) = *N*_*features*_(*cortical surface area*) = 5124). Volumetric data for subcortical regions and measures of global volume were extracted from the aseg.stats file (*N*_*features*_(*subcortical*) = 66).

#### 3.1.2. Predictive analysis

Predictive models were implemented in a two-level approach (see Figure 1.4). On the first level, linear support vector regression models (SVR, Drucker et al., 1996) were used to predict age from neuroimaging data (single-source models). On the second level, predictions from the single-source models were stacked with random forest (RF, Breiman, 2001) regression models. Using RF models for stacking multiple neuroimaging modalities has previously been shown to produce better predictions with smaller variability in prediction errors as compared to other stacking methods (Rahim et al., 2016). All predictive analyses have been performed using the python-based Scikitlearn package (Pedregosa et al., 2011). The code is available at github.com/fliem/LeiCA_LIFE.

In detail, this procedure entailed (see Figure 1):

1. *Train-test-split*. Data was split into equally sized training and test set (see Figure 1.3). The training set was used to train (learn) the models, the test set was put aside to subsequently evaluate the models’ performance on unseen data.
2. *Training of single-source models (see Figure 1.4)*. Two parallel approaches have been used to train single-source models. First, using the neuroimaging data of the entire training set, single-source SVR models were fitted, resulting in one trained model per source. Second, in parallel, using the neuroimaging data of the training set, single-source SVR models were trained in a 5-fold cross-validation (CV) approach (see Figure 1.4.1). This was done to obtain unbiased CV-predictions (to be used in the following step).
3. *Training of multi-source models*. To aggregate information from multiple sources into one prediction, the previous step’s CV-predictions were stacked (concatenated). A feature vector based on the single-source predictions was constructed. Based on this feature vector, the multi-source models were fitted, to obtain trained multi-source models (see Figure 1.4.2). This was done in three versions:
  (a) *stacked-function* which combined age predictions from *connectivity matrix 197* and *connectivity matrix 444,*
  (b) *stacked-anatomy* which combined age predictions from *cortical thickness, cortical surface area, and subcortical volumes,* and
  (c) *stacked-multimodal* which combined age predictions from all five single-source models. For instance, in the case of *stacked-multimodal,* each subject’s feature vector consisted of the five age prediction values from the single-source models.
4. *Test the models by predicting age in new subjects*. The performance of the trained single- and multi-source models was tested with the neuroimaging data of the test set as input (see Figure 1.5). First, single-source predictions are calculated by using the trained model from step 2 (see Figure 1.5.1). Second, these predictions are stacked and fed into the trained model from step 3 (see Figure 1.5.2), to receive single- and multi-source test set predictions.
5. *Evaluate generalization performance*. Finally, the models’ generalization performance can be assessed via the test set’s absolute error (AE) of the age predictions (obtained in step 4) from chronological age. Additionally, the coefficient of determination (R2) is also reported.

*Statistical tests*. To compare models, non-parametric statistical tests were run on the absolute prediction errors, using the SciPy package (v0.17.0, scipy.org). Correction for multiple comparisons was done using the false discovery rate (FDR) procedure described by Benjamini & Hochberg (1995). Results were plotted with the Seaborn package (v0.7.0, Waskom et al., 2016).

*Tuning of hyperparameters*. For the single-source SVR models, tuning curves for the *C* parameter were run on the training data. These curves showed a ‘sweet spot’ for the high dimensional neuroimaging input data (the connectivity matrices, cortical thickness and cortical surface area) around *C* = 1e–3 (for an example see Figure A.6). For the lower dimensional subcortical input data, the standard *C* = 1 performed well. All models were run with the default *ε* = 0.1. For the multi-source RF models, out-of-bag estimates were used to set the tree depth.

### 3.2. Analysis plan

A brief sketch of the different analyzes, tailored to the different research questions, follows here. Further details can be found in the results section.

#### 3.2.1. Prediction performance in multimodal data

Age predictions have been performed as described in section 3.1. The entire LIFE sample was split into equally sized training and test set.

#### 3.2.2. Brain aging in cognitively impaired subjects

For this analysis, the sample was reduced to subjects with a valid OCI score (see section 2.2). Models were trained on *OCI-norm* subjects only. The test set consisted of subjects from all OCI groups. A brain aging (BA) score was then calculated for each subject and each single- and multi-source model by subtracting *age*_*chronological*_ from *age*_*predicted*_. These BA scores were compared between OCI groups.

#### 3.2.3. Robustness against confounds

*Head motion*. The robustness of our approach against head motion was investigated with the models described in section 3.2.1. First, we did this using *motion regression*. On the group-level, head motion (mean FD derived from the functional scans) was regressed out of the input matrices in the single-source models for the entire training and entire test set separately. Mean FD was derived from the functional acquisition and used as a measure of head motion for the functional as well as the structural data. Second, as an alternative, *motion matching* was performed by creating a subsample of the test set that does not show an *age* × *motion* correlation. For direct comparison, an equally sized random sample was also drawn. These test samples were then used to evaluate the performance of the models trained on the full training sample (see 3.2.1).

*Generalization to new site*. In this step we investigated how the models generalize to data from a new site (different country, scanner, acquisition protocol, subjects). Age predictions were performed on data from the NKI set using models trained on LIFE data (one sample training; section 3.2.1). In a subsequent analysis, models were re-trained on a training set that combined the original LIFE training sample with a small number of subjects from the NKI set, to increase the generalizability of the predictive models (two sample training). Finally, to increase the training sample size, the one and two sample training was repeated, including the majority of the LIFE data into the training sample, not only the original training set (99 % of the entire LIFE sample; 1% was retained for a rough check of the models on LIFE data). This analysis will be referred to as *full LIFE sample training* approach.

## 4. Results

Our results demonstrate that i) incorporating multiple brain imaging modalities increases age prediction performance (Figure 2 and 3); ii) subjects with objective cognitive impairment show advanced brain aging compared to subjects without objective cognitive impairment (Figure 4); iii) our prediction models are robust against confounds (Figure 5), i.e., not driven by head motion and generalize to new datasets. For the comparison of modalities, data for models of all modalities will be presented. After showing that the multimodal approach outperforms the others, the remainder of the results, for the sake of brevity, will focus on this model. The full results can be found in Appendix B.

**Figure 2.**
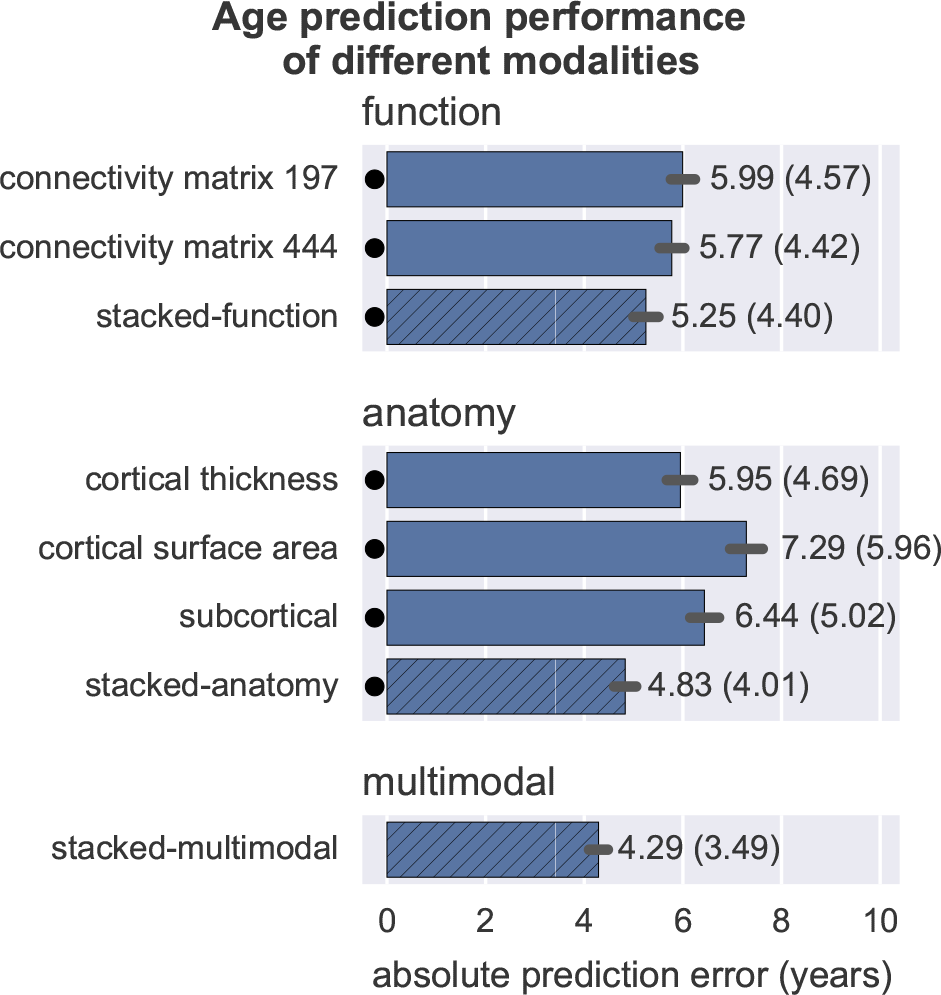
Prediction performance (absolute error on test set, lower values are better). Note that the *stacked-multimodal* model shows the least prediction error. Black dots represent significant (*p*(*FDR*) < 0.05) deviation from the best model, indicating that *stacked-multimodal* significantly outperforms all other models. Error bars represent 95% CI bootstrapped with 1000 iterations. Numbers next to error bars represent mean (standard deviation). Stacked models are shown with hatched bars. For full statistics see Table B.1

**Figure 3.**
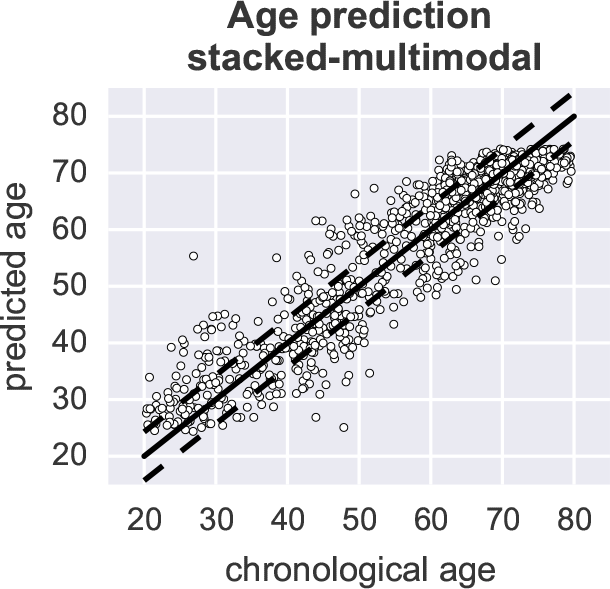
Chronological and predicted age from the *stacked-multimodal* model. Circles represent subjects, the solid line the perfect prediction, dashed lines the mean absolute prediction error (4.29 years).

**Figure 4.**
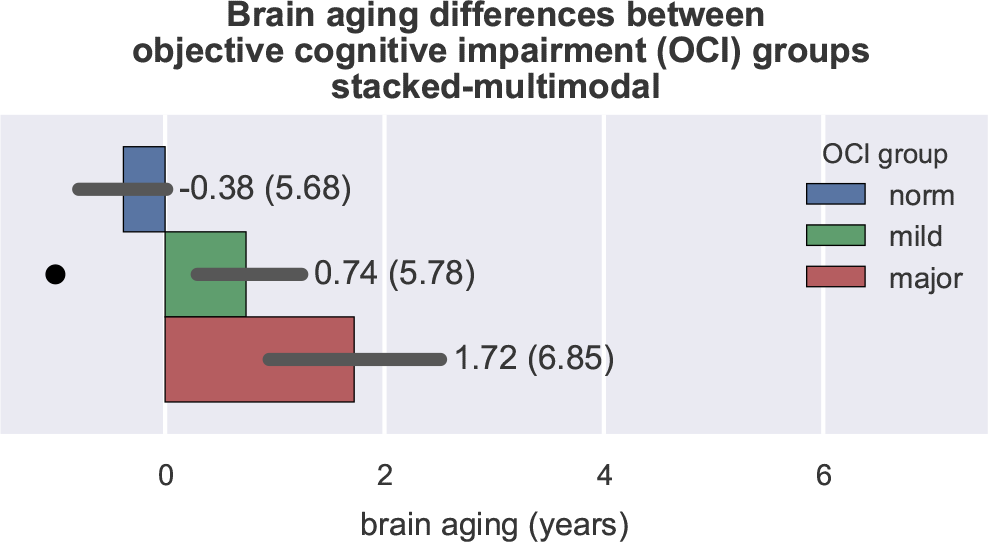
Differences in brain aging (brain aging = predicted age - chronological age) between OCI groups for *stacked-multimodal*. Positive brain aging values indicate that a brain appears older than expected from chronological age. Note that brain aging significantly increases with severity of OCI, i.e., more advanced brain aging in OCI. Numbers next to error bars represent mean and standard deviation. For full data see Figure B.7 and Table B.6.

**Figure 5.**
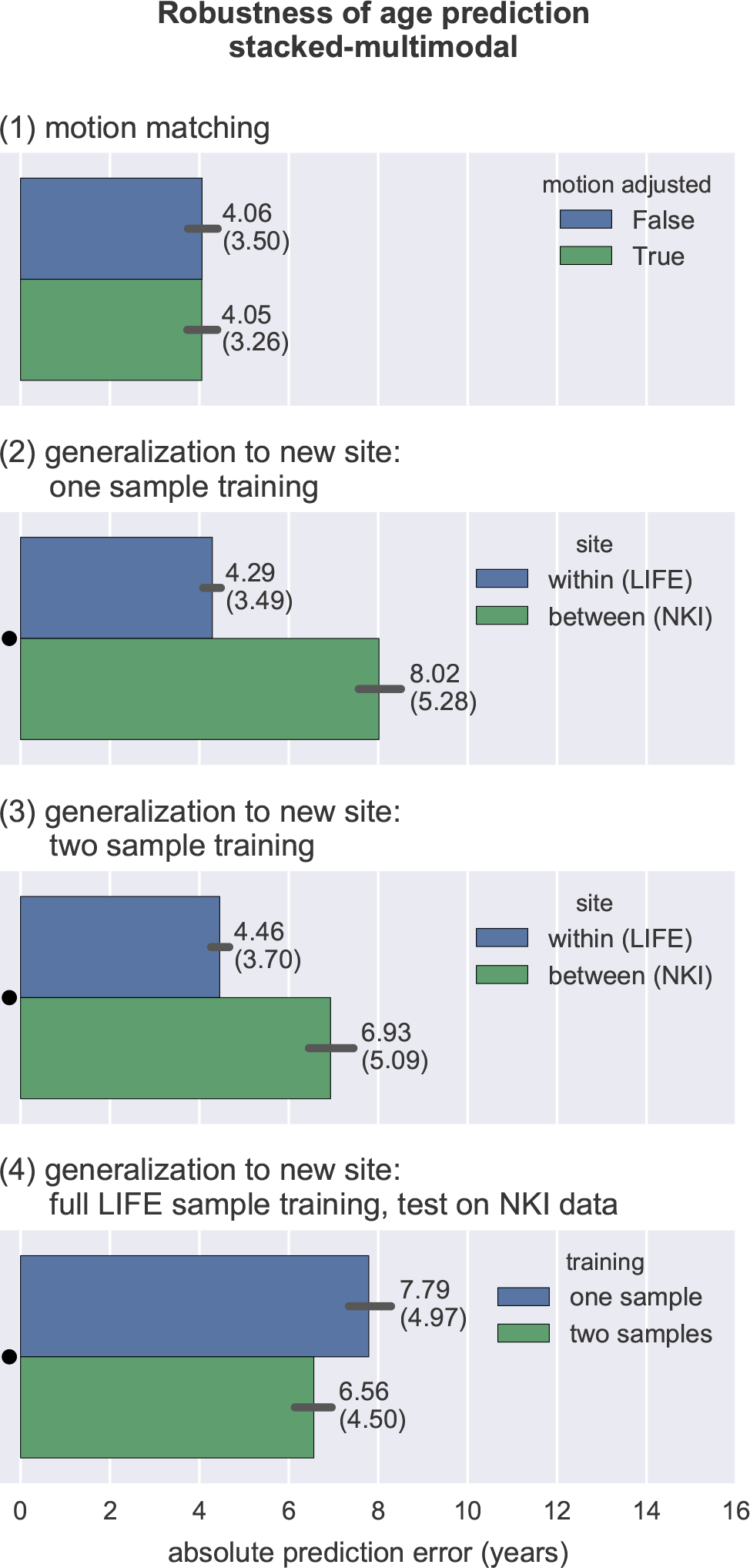
Robustness of age prediction against confounds for the *stackedmultimodal* model. (1) Motion matching analysis show that age prediction works equally good in motion adjusted (without *age* × *motion* correlation) and non-adjusted (with *age* × *motion* correlation) groups, indicating that the predictive model is not driven by head motion. Full data are shown in Figure B.9 and Table B.8. Note that the slightly lower prediction error (around 4.06) as compared to the original analysis (around 4.29; see Figure 2) is a result of the restricted age range of the test samples in the motion matching analysis. Hence, those values should only be compared within the motion matching analysis and not with the original analysis. (2) Generalization to new site. Standard training procedure (one sample training) showed significantly (*p*(*FDR*) < 0.05 as indicated by the black dot) better prediction performance in LIFE data (within site) than in NKI data (between site, for full data see Figure B.10 and Table B.9). (3) After training the model on a mixed-site sample (two sample training, *N*_*LIFE*_ = 1177; *N*_*NKI*_ = 46), predictions on the NKI data improve (Table B.10), but the predictions on the main training site LIFE (within site) still are significantly better than on the minor training site NKI (between site, for full data see Figure B.11 and Table B.11). (4) Finally, generalization is investigated by training on the full LIFE sample (*N*_*training,LIFE*_ = 2377). Test prediction performance on between-site NKI data for one sample training (LIFE sample only) and two samples (LIFE + NKI samples; *N*_*training,NKI*_ = 46) slightly increases as compared to the original training approach (green bars from (2) and (3); for full data see Figure B.12 and Table B.12).

### 4.1. Multimodal data increases prediction performance

Based on the entire LIFE sample, age prediction models were trained on the training set (*N* = 1177) and evaluated on the test set (*N* = 1177). Figure 2 shows prediction performance on the test sample (for full statistics see Table B.1). All models show good prediction performance (mean absolute error between 4.29 and 7.29 years, R2 between 0.62 and 0.87). The stacked models show better performance than single source models, with the *stacked-multimodal* model outperforming all other models. Additionally, this model also shows the least prediction variability. By going from the second best model, *stacked-anatomy,* to the best, *stacked-multimodal,* approximately half a year in prediction accuracy is gained. Table B.2 shows feature importances for the multi-source models.

Figure 3 shows the individual predictions for the *stacked-multimodal* model, the model with the best predictive performance.

### 4.2. Advanced brain aging in cognitively impaired subjects

Based on a large battery of cognitive tests, subjects with mild or major OCI were identified. For this analysis, the models were trained on half of the *OCI-norm* subjects (*N*_*training*_ = 724). Subsequently, age predictions were performed on a test sample containing *OCI-norm, mild* and *major* subjects (test: *Ν _norm_* = 729, *N*_*mild*_ = 632, *N*_*major*_ = 251) and compared between groups (sample characteristics can be found in Table B.4). Figure 4 shows the advanced brain aging in OCI predicted by the *stacked-multimodal* model. Figure B.7 shows that all models, except *stacked-function,* show a significant progression in brain aging related to the severity of OCI (see Table B.6 for full statistics). This finding demonstrates that the age prediction models capture aspects of cognitive impairment.

### 4.3. Robustness against confounds

*Head motion*. Our sample showed a substantial *age* × *motion* correlation (*r*_*age*×*motion*_ = 0.43; *p* = 1.87e–15; head motion defined as mean FD derived from the functional scans). To test the influence of head motion on the age prediction models, the following two analyses have been performed.

*Motion regression*. To test the models’ robustness against head motion, we regressed out head motion (mean FD) from the input data (for training and test set separately). Regressing out motion reduces prediction accuracy significantly (e.g., the *stacked-multimodal* model’s error increases from 4.29 to 6.95 years; see Figure B.8 and Table B.7). This might either be the result of head motion driving the prediction models, or, due to the large variance shared by age, head motion and the brain measures, of too aggressively removing age-related variance while removing motion-related variance. To test these two alternative explanations, the following motion matching analysis was performed.

*Motion matching*. In this analysis, a motion adjusted subsample of the test set is created by restricting the sample to subjects with a mean FD between 0.19 and 0.28 mm and an age above 25 years, which results in a motion matched subsample (*N* = 387; *r*_*age*×*motion*_ = 0.06; *p* = 0.26). Excluding subjects with a mean FD lower than 0.19 was necessary to create a balanced sample, because of the dominance of young subjects with low motion. An equally sized non-motion-adjusted subsample was randomly drawn for comparison (*r*_*age*×*motion*_ = 0.42; *p* = 2.45e–18). These subsamples (*N* = 387) were used to evaluate the influence of motion in the models trained on the original training set (*N* = 1177; see 4.1). The *stacked-multimodal* model (Figure 5.1), as well as all other models (Figure B.9) perform equally well with and without motion matching (all *p* > 0.49; for full statistics see Table B.8), indicating that head motion is not driving the age prediction models and that motion regression is removing too much meaningful age-related variance.

*Generalization to new site*. To demonstrate how the models generalize to data from a new site (different country, scanner, acquisition protocol, subjects), we predicted age on NKI data with models that have been trained on LIFE data (one sample training). While the models perform much better than chance, unsurprisingly, better predictive performance is achieved on LIFE than on NKI data (Figure 5.2; for full data see Figure B.10 and Table B.9). Assuming that models show higher generalizabilty if trained on more heterogeneous data, the following post-hoc analysis tested whether adding a small number of subjects from NKI to the LIFE training sample increases generalization (two sample training; training sample: *N*_*LIFE*_ = 1177; *N*_*NKI*_ = 46, representing around 10% of the NKI sample).

While prediction performance increases by adding subjects from NKI to the training sample (see Table B.10), Figure 5.3 shows that prediction still works better on LIFE data (for full data see Figure B.11 and Table B.11). We also demonstrate that these results are robust across different random splits of the data (see Table B.13).

As a further attempt to increase generalizability, we pursued the *full LIFE sample training* approach. Here, we repeated the one and two sample training using the majority of all LIFE data for training (training samples: one sample training: *N*_*LIFE*_ = 2377); two sample training. *N*_*LIFE*_ = 2377; *N*_*NKI*_ = 46). These trained models were then evaluated with the (remaining) NKI data. This further increases generalizability and reduced the prediction error. For the *stacked-multimodal* (two sample) analysis it decreases to 6.56 years (Figure 5.4), which is a slight reduction compared to the original two sample result of 6.93 years (for full data see Table B.12 and Figure B.12, for a scatter plot of test predictions Figure B.13).

## 5. Discussion

The aim of the current study was to establish a novel multimodal brain-based age prediction framework that makes use of information from anatomy and functional connectivity. We found that (i) including multimodal information increases prediction accuracy, (ii) objective cognitive impairment is associated with increased brain aging, and (iii) our framework is robust against confounds, most importantly, against head motion, and generalizes to new datasets, especially if the training set is composed of a large and heterogeneous dataset.

Age prediction was best achieved using the multimodal approach (*stacked-multimodal*), which showed a mean absolute age prediction error of 4.29 years. This is approximately a half-year more accurate than when only taking anatomical information into account (*stacked-anatomy*). Furthermore, the multimodal approach shows less variability in prediction performance. We assume that the gain in prediction accuracy is a result of the different brain-imaging modalities’ shared variance, via reducing the measurement error of brain data, as well as unique variance, via the addition of new information. Aggregating multiple sources of neuroimaging data via Random Forest models has been shown to work well (Rahim et al., 2016). In particular, aggregating data via RF models results in better age prediction performance as compared to merely averaging single-source predictions (e.g., for *stacked-multimodal:* age prediction error of 4.29 vs 5.08 years).

Our anatomical approach is conceptually similar to the framework of Franke et al. (2010). The main difference is the choice of anatomical data analysis tool: voxel-based morphometry in their work, surface-based morphometry in ours. Their best model showed a mean absolute prediction error of 4.61, which is in agreement with the performance of *stacked-anatomy* at 4.83 years. The surface-based morphometry approach has the advantage of disentangling structural information of cortical thickness and surface area (Meyer et al., 2014). Age prediction based on cortical thickness worked better than based on cortical surface area, which is well in line with stronger age-related effects in cortical thickness than in surface area (Hogstrom et al., 2013). Future studies might also investigate whether considering additional information about white matter anatomy further reduces prediction accuracy. How much further the prediction accuracy can be reduced, i.e., the lower bound, is unclear. Due to individual differences in the brains of individuals of the same age, some prediction error will always persist.

We investigated brain aging in individuals with objective cognitive impairment. By subtracting chronological age from predicted age, we calculated a brain aging score (also called *brainAGE (brain age gap estimate)* by Franke et al. (2010), or *PAD (predicted age difference)* by Cole et al. (2015)). The multimodal approach, as well as most other approaches we investigated, predicted significantly increased brain aging in participants with objective cognitive impairment. The progression of brain aging always followed the progression of OCI and increased from normal to mild to major OCI individuals. The strongest differences in brain aging between the OCI groups was observed in the model using subcortical data. This suggests that while the multimodal approach performed best in age prediction, differences in cognitive performance might be better characterized using specific modalities. As different pathologies might be detectable early in different MRI modalities, future studies should consider the effectiveness of predictive models of different uniand multimodal approaches in the context of a given pathology.

The brain-age metric provides an interpretable aggregate measure of brain aging processes in brain structure and function. However, if the primary research interest is predicting cognitive performance, why investigate this via metrics of brain-age? Ideally, the predictive model should be created using a study-specific cognitive target (for instance, see Ullman et al., 2014). Directly predicting future cognitive performance certainly holds tremendous potential to identify specific cognitive modalities at risk of future decline. These models offer a valuable foundation for innovating tailored interventions through, for example, cognitive training.

However, to obtain stable models large datasets with brain and behavioral data is required. Assessment of brain-age offers an alternative and complementary measure that is already available through several publicly available large-scale brainimaging datasets. Such data can be used to train models, which are then complemented by a smaller, but richer dataset that includes information about cognitive performance, in order to test specific hypotheses.

Confounding effects of head motion in brain-imaging studies have received increased interest in the recent years (e.g., see Power et al., 2012; Reuter et al., 2015). The present study demonstrated that head motion does not drive brain-based age prediction and that regressing out motion might also affect meaningful age-related variance. While the estimation of head motion in functional MRI scans is well established, this is much more challenging for structural MRI scans due to their longer acquisition times. While there exist special acquisition protocols tailored to measure head motion (for instance see Reuter et al., 2015), these are not yet standard, and do not apply to already existing data. However, since head motion has within-subject stability (Van Dijk et al., 2012), we took head motion estimates based on functional scans as a proxy for head motion in structural scans, as also done by Alexander-Bloch et al. (2016). Nevertheless, motion between different scan blocks certainly can differ, which might render the motion metrics derived from functional scans a poor proxy for structural scans. For instance, the time point of collecting the structural data (at the beginning or end of a scanning session) might result in different motion characteristics due to fatigue of the study participants or adaptation to the in-scanner situation. These effects have not yet been studied systematically and deserve attention in future studies.

Age prediction models perform significantly better when trained and tested on data from the same site, as compared to data from different sites. Training models on a larger and more heterogeneous dataset modestly improves the prediction accuracy. However, since even in this case within-site prediction outperforms between-site prediction this topic deserves more attention in future work. Several factors may have contributed to the better generalizability observed in the NKI dataset using anatomical rather than functional information. First, the anatomical sequences used in both studies are quite similar, while the functional sequences differ with regards to temporal and spatial parameters. Second, anatomical information analyzed with surface-based morphometry shows higher reliability (Liem et al., 2015) than functional MRI (Shehzad et al., 2009). To avoid fitting models to the idiosyncrasies of a given study, future studies should broaden the variability of training data by including data from an array of sites, as recently demonstrated (Abraham et al., 2016; Cole et al., 2015).

A standardization of MRI acquisition protocols may also contribute to a better generalization of predictive models. Quantitative structural MRI (Lutti et al., 2014) or calibration of functional sequences (Chiarelli et al., 2007) may provide more reliable and valid brain measurements, resulting in better predictors. However, these techniques are not currently standard practice and require further development for widespread application (e.g., see Dubois & Adolphs, 2016).

By moving from correlative studies to predictive studies using tools from machine learning (Gabrieli et al., 2015; Yarkoni & Westfall, 2016), cognitive neuroscience as a basic science might be complemented with an applied component that can give relevant insights into both clinical pathologies as well as the healthy spectrum of aging. This may range from brain-based biomarkers for neurological or psychiatric diseases, to identifying potential future cognitive impairments on an individual-level and designing targeted cognitive training.

## 6. Conclusions

In the present study, we demonstrated that including information from multiple MR modalities, i.e., anatomy and functional connectivity, increased accuracy of brain-based age prediction. Brain-age measured with this multimodal framework was accelerated in subjects with cognitive impairment. Importantly, head motion does not drive brain-based age prediction and predictive models generalize to new datasets, especially if those are trained on large and heterogeneous datasets. Given these findings, measuring brain aging using machine learning methods holds promise for establishing brain-based biomarkers that could aid diagnosis of neurocognitive disorders and be relevant for clinical practice.

## 7. Acknowledgments

The first author thanks all colleagues inside and outside the Max Planck Institute for Human Cognitive and Brain Sciences that provided valuable feedback for this project, especially the members of the Neuroanatomy and Connectivity Group.

We would like to thank the Enhanced Nathan Kline Institute-Rockland Sample initiative for sharing their data.

Franziskus Liem is supported by the Swiss National Science Foundation (SNSF), grant number P2ZHP1_155200. Gael Varoquaux and Mehdi Rahim are supported by the NiConnect project (ANR-11-BINF-0004 NiConnect). Jana Kynast is supported by the Max-Planck International Research Network on Aging (MaxNetAging).

This work is supported by the European Union, the European Regional Development Fund, and the Free State of Saxony within the framework of the excellence initiative, and LIFE-Leipzig Research Center for Civilization Diseases, University of Leipzig (project numbers 713-241202, 713-241202, 14505/2470, 14575/2470), and by the German Research Foundation (CRC1052 Obesity mechanisms Project A01).

## Appendix A. Supplementary methods

### Appendix A.1. Tuning curve

**Figure A.6:**
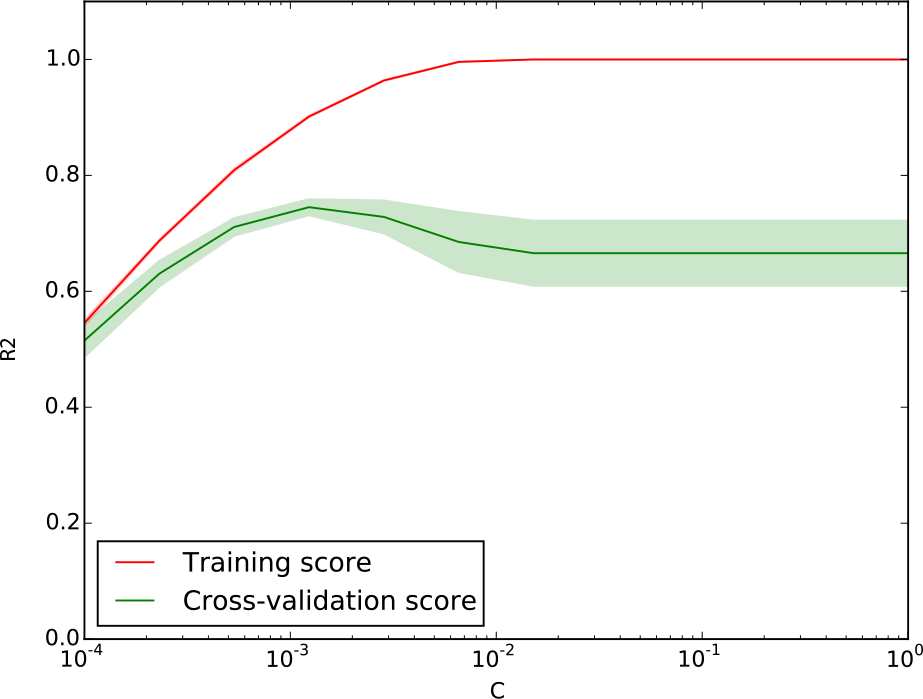
Exemplarily, a tuning curve is presented for *cortical thickness*. The tuning curve was run on the training set. The support vector regression’s *C* parameter (x-axis) was varied, the R2 (coefficient of determination, y-axis) was evaluated as training and cross-validation score. A ‘sweet spot’, a maximum cross-validation score (green line) only a slightly better training performance (red line), can be seen around *C* = 10^−3^.

### Appendix A.2. NKI sample

Data from the enhanced Nathan Kline Institute – Rockland sample (fcon_1000.projects.nitrc.org/indi/enhanced, Nooner et al., 2012) was used in parts of this study. Subjects selected for the present work (N = 475) were between 18 and 85 years (M = 45.78; SD = 18.91; 311 female, 161 male).

### Appendix A.3. MR data

Brain imaging was performed on a a 3T Siemens Trio scanner with a 32 channel head coil.

T2*-weighted functional images were acquired using an multiband echo-planar-imaging sequence with 3 mm isotropic voxels, 40 slices, echo time (TE) of 30 ms, repetition time (TR) of 645 ms, multiband acceleration factor of 4 and a flip-angle of 60° (fcon_1000.projects.nitrc.org/indi/enhanced/mri_protocol.html). The resting-state sequence lasted approximately 9.5 min (900 volumes), during which subjects were instructed to keep their eyes open and not to fall asleep. No fieldmaps have been acquired.

High resolution T1-weighted structural images were acquired using the MP-RAGE sequence with 1 mm isotropic voxels, 176 slices, a TR of 1900 ms, and a TE of 2.52 ms.

### Appendix A.4. MR data preprocessing

Preprocessing was performed very similar to preprocessing of the LIFE sample detailed in section 2.4. Since no fieldmaps were available for NKI and data from the two studies have been preprocessed independently, minor differences exist: CompCor was performed with five components instead of six and normalization into standard space was performed with FSL’s FNIRT, not ANTS. Preprocessing scripts are available at github.com/fliem/nki_nilearn.

## Appendix B. Supplementary results

### Appendix B.1. Multimodal data increases age prediction performance

**Table B.1:** Age prediction (absolute prediction error (APE)) on the test sample. Statistical test against best model (Wilcoxon signed-rank test against stacked-multimodal; N = 1177). See Figure 2.

|  | M(APE) | SD(APE) | R2 | W | p(FDR) |
| --- | --- | --- | --- | --- | --- |
| connectivity matrix 197 | 5.99 | 4.57 | 0.75 | 2e+05 | 8.2e-38 |
| connectivity matrix 444 | 5.77 | 4.42 | 0.77 | 2.1e+05 | 9.1e-31 |
| stacked-function | 5.25 | 4.40 | 0.80 | 2.3e+05 | 1.6e-22 |
| cortical thickness | 5.95 | 4.69 | 0.75 | 2.1e+05 | 9.9e-32 |
| cortical surface area | 7.29 | 5.95 | 0.62 | 1.7e+05 | 5.9e-53 |
| subcortical | 6.44 | 5.02 | 0.71 | 1.9e+05 | 1.6e-40 |
| stacked-anatomy | 4.83 | 4.01 | 0.83 | 2.7e+05 | 1.8e-10 |
| stacked-multimodal | 4.29 | 3.49 | 0.87 | - | - |

### Appendix B.2. RF feature importance

**Table B.2:**
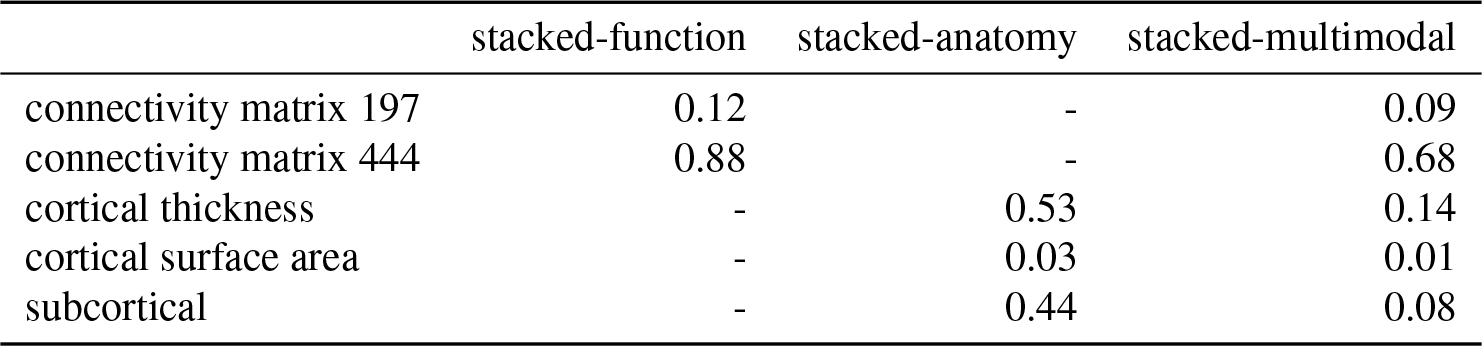
Values of feature importance from the Random Forest multi-source models showing the contribution of the single-source data.

|  | stacked-function | stacked-anatomy | stacked-multimodal |
| --- | --- | --- | --- |
| connectivity matrix 197 | 0.12 | - | 0.09 |
| connectivity matrix 444 | 0.88 | - | 0.68 |
| cortical thickness | - | 0.53 | 0.14 |
| cortical surface area | - | 0.03 | 0.01 |
| subcortical | - | 0.44 | 0.08 |

### Appendix B.3. Cross-validation (CV) scores

**Table B.3:** CV scores of five folds run on training sample.

|  | M(APE) | SD(APE) | M(R2) | SD(R2) |
| --- | --- | --- | --- | --- |
| connectivity matrix 197 | 6.33 | 0.31 | 0.72 | 0.03 |
| connectivity matrix 444 | 6.02 | 0.40 | 0.75 | 0.03 |
| stacked-function | 5.45 | 0.20 | 0.78 | 0.02 |
| cortical thickness | 6.25 | 0.12 | 0.73 | 0.02 |
| cortical surface area | 7.21 | 0.34 | 0.61 | 0.04 |
| subcortical | 6.60 | 0.34 | 0.69 | 0.04 |
| stacked-anatomy | 4.98 | 0.10 | 0.82 | 0.02 |
| stacked-multimodal | 4.53 | 0.12 | 0.85 | 0.01 |

### Appendix B.4. Increased brain aging in impaired subjects

**Table B.4:** Sample characteristics of OCI (objective cognitive impairment) groups of the test sample.

| OCI group | N | M(Age) | SD(Age) | Sex(f/m) |
| --- | --- | --- | --- | --- |
| norm | 729 | 59.2 | 15.2 | 364/365 |
| mild | 632 | 58.0 | 14.9 | 294/338 |
| major | 251 | 58.3 | 15.7 | 115/136 |

**Table B.5:**
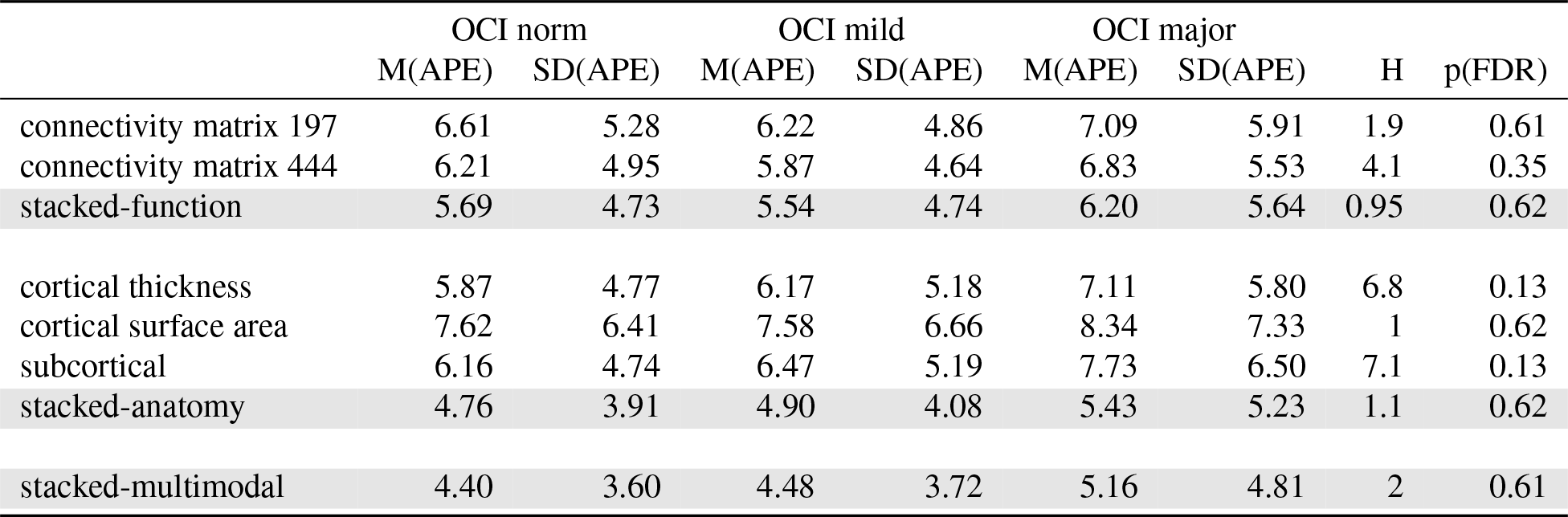
Differences in absolute prediction error (APE) between objective cognitive impairment (OCI) groups (Kruskal-Wallis H-test: effect of OCI group (norm, mild, major) on absolute prediction error (APE); N(norm) = 729, N(mild) = 632, N(major) = 251; N(training) = 724.

|  | OCI norm |  | OCI mild |  | OCI major |  | H | p(FDR) |
| --- | --- | --- | --- | --- | --- | --- | --- | --- |
|  | M(APE) | SD(APE) | M(APE) | SD(APE) | M(APE) | SD(APE) |  |  |
| connectivity matrix 197 | 6.61 | 5.28 | 6.22 | 4.86 | 7.09 | 5.91 | 1.9 | 0.61 |
| connectivity matrix 444 | 6.21 | 4.95 | 5.87 | 4.64 | 6.83 | 5.53 | 4.1 | 0.35 |
| stacked-function | 5.69 | 4.73 | 5.54 | 4.74 | 6.20 | 5.64 | 0.95 | 0.62 |
| cortical thickness | 5.87 | 4.77 | 6.17 | 5.18 | 7.11 | 5.80 | 6.8 | 0.13 |
| cortical surface area | 7.62 | 6.41 | 7.58 | 6.66 | 8.34 | 7.33 | 1 | 0.62 |
| subcortical | 6.16 | 4.74 | 6.47 | 5.19 | 7.73 | 6.50 | 7.1 | 0.13 |
| stacked-anatomy | 4.76 | 3.91 | 4.90 | 4.08 | 5.43 | 5.23 | 1.1 | 0.62 |
| stacked-multimodal | 4.40 | 3.60 | 4.48 | 3.72 | 5.16 | 4.81 | 2 | 0.61 |

**Table B.6:** Differences in brain aging (BA) between objective cognitive impairment (OCI) groups (Kruskal-Wallis H-test: effect of OCI group (norm, mild, major) on brain age; N(norm) = 729, N(mild) = 632, N(major) = 251). N(training) = 724. See Figure B.7.

|  | OCI norm |  | OCI mild |  | OCI major |  | H | p(FDR) |
| --- | --- | --- | --- | --- | --- | --- | --- | --- |
|  | M(BA) | SD(BA) | M(BA) | SD(BA) | M(BA) | SD(BA) |  |  |
| connectivity matrix 197 | 0.48 | 8.45 | 1.31 | 7.79 | 2.22 | 8.97 | 7.9 | 0.026 |
| connectivity matrix 444 | -0.04 | 7.95 | 0.75 | 7.45 | 1.52 | 8.66 | 7.5 | 0.027 |
| stacked-function | -0.24 | 7.40 | 0.64 | 7.26 | 1.22 | 8.30 | 5.1 | 0.077 |
| cortical thickness | 0.73 | 7.54 | 2.16 | 7.76 | 2.99 | 8.68 | 21 | 5.1e-05 |
| cortical surface area | 1.32 | 9.87 | 2.82 | 9.69 | 3.31 | 10.61 | 15 | 0.001 |
| subcortical | -0.12 | 7.78 | 1.39 | 8.18 | 3.91 | 9.32 | 44 | 2.1e-09 |
| stacked-anatomy | -0.52 | 6.14 | 0.76 | 6.33 | 1.88 | 7.31 | 26 | 7.8e-06 |
| stacked-multimodal | -0.38 | 5.68 | 0.74 | 5.78 | 1.72 | 6.85 | 22 | 4e-05 |

**Figure B.7:**
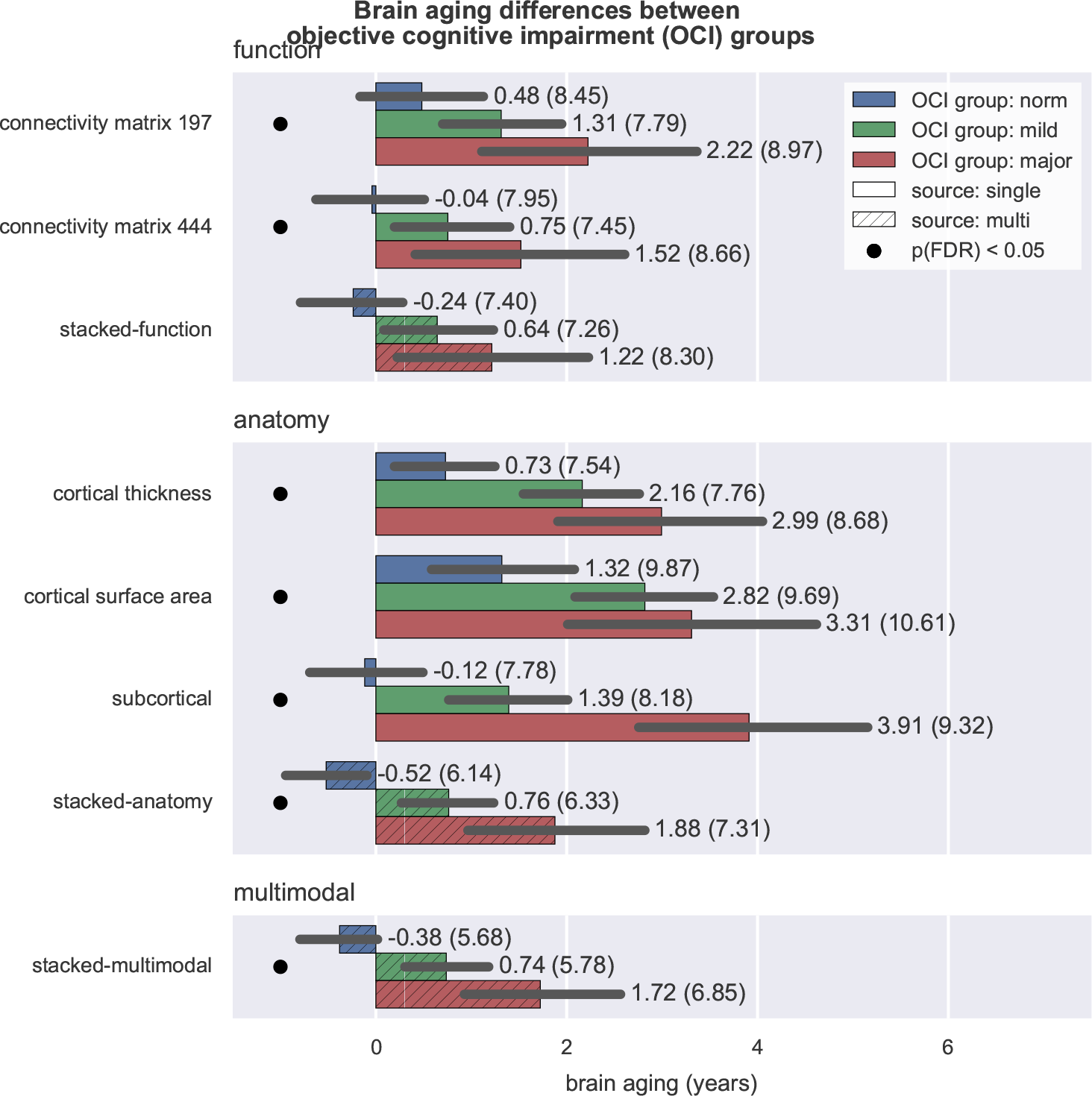
Differences in brain aging between objective cognitive impairment (OCI) groups. Positive brain aging values indicate that a brain appears older than expected from chronological age. Note that for the majority of models impairment measured by OCI is significantly related to higher brain aging (as indicated by the black dot), i.e., more advanced brain aging. For full statistics see Table B.6.

### Appendix B.5. Robustness against head motion

#### Appendix B.5.1. Motion regression

**Figure B.8:**
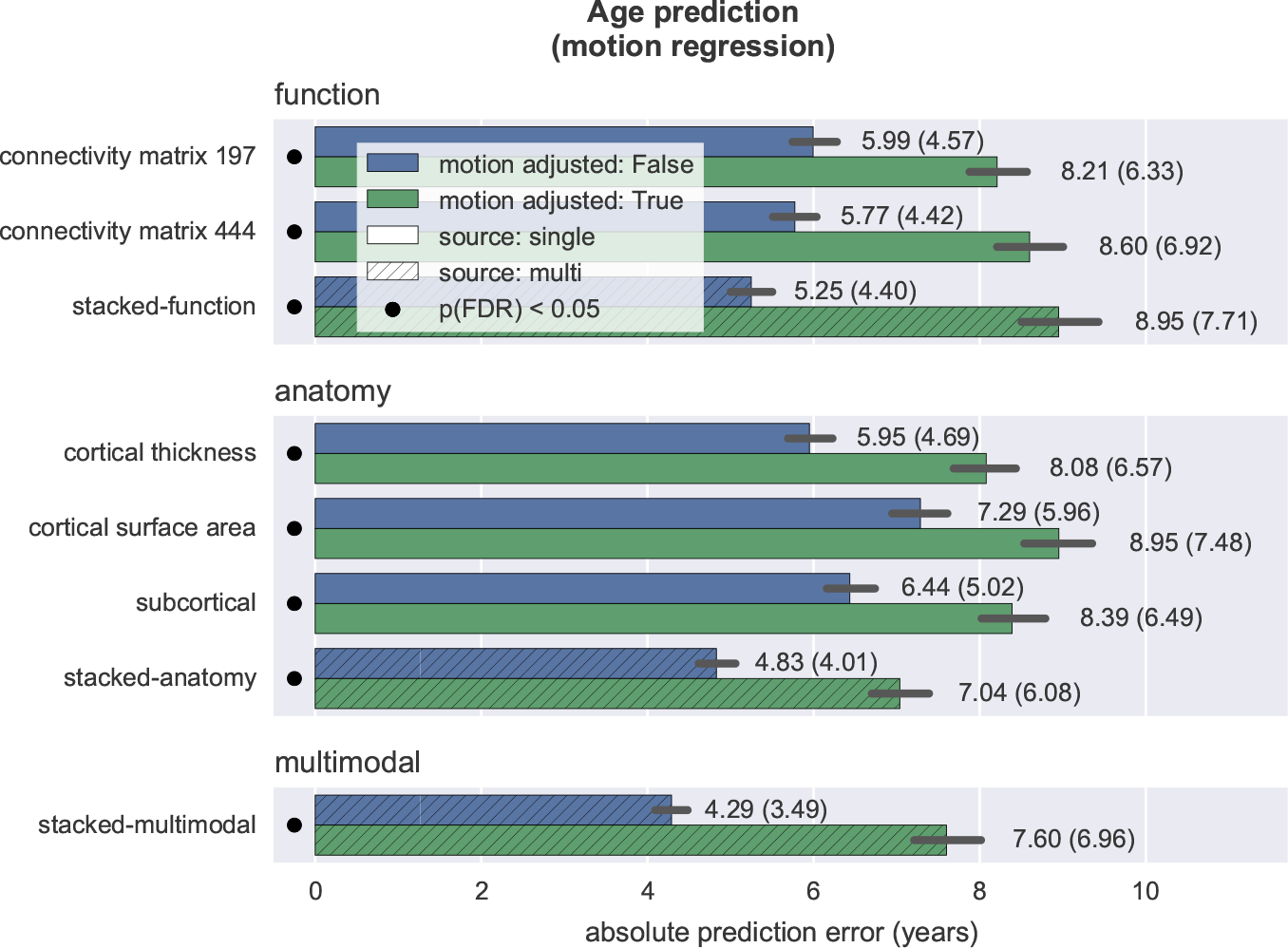
Motion adjustment via motion regression. Absolute prediction error significantly increases if motion is regressed out of brain data (black dots). Motion adjusted: True: analysis includes motion regression on brain data; False: original analysis without motion regression.

**Table B.7:** Motion adjustment via motion regression. Absolute prediction error (APE) significantly increases if motion is regressed out of brain data. Motion adjusted: True: analysis includes motion regression on brain data; False: original analysis without motion regression. R2: coefficient of determination. (Wilcoxon signed-rank test: motion adjusted False against True; Ν = 1177). See Figure B.8.

|  | motion adjusted False |  |  | motion adjusted True |  |  | W | p(FDR) |
| --- | --- | --- | --- | --- | --- | --- | --- | --- |
|  | M(APE) | SD(APE) | R2 | M(APE) | SD(APE) | R2 |  |  |
| connectivity matrix 197 | 5.99 | 4.57 | 0.75 | 8.21 | 6.33 | 0.53 | 1.9e+05 | 1.3e-39 |
| connectivity matrix 444 | 5.77 | 4.42 | 0.77 | 8.60 | 6.92 | 0.47 | 1.8e+05 | 1.7e-46 |
| stacked-function | 5.25 | 4.40 | 0.80 | 8.95 | 7.71 | 0.40 | 1.7e+05 | 3.2e-48 |
| cortical thickness | 5.95 | 4.69 | 0.75 | 8.08 | 6.57 | 0.53 | 1.9e+05 | 1e-42 |
| cortical surface area | 7.29 | 5.95 | 0.62 | 8.95 | 7.48 | 0.41 | 2.1e+05 | 3.6e-33 |
| subcortical | 6.44 | 5.02 | 0.71 | 8.39 | 6.48 | 0.51 | 2.1e+05 | 2e-33 |
| stacked-anatomy | 4.83 | 4.01 | 0.83 | 7.04 | 6.08 | 0.63 | 2.1e+05 | 3.6e-33 |
| stacked-multimodal | 4.29 | 3.49 | 0.87 | 7.60 | 6.95 | 0.54 | 1.8e+05 | 7.3e-47 |

#### Appendix B.5.2. Motion matching

**Figure B.9:**
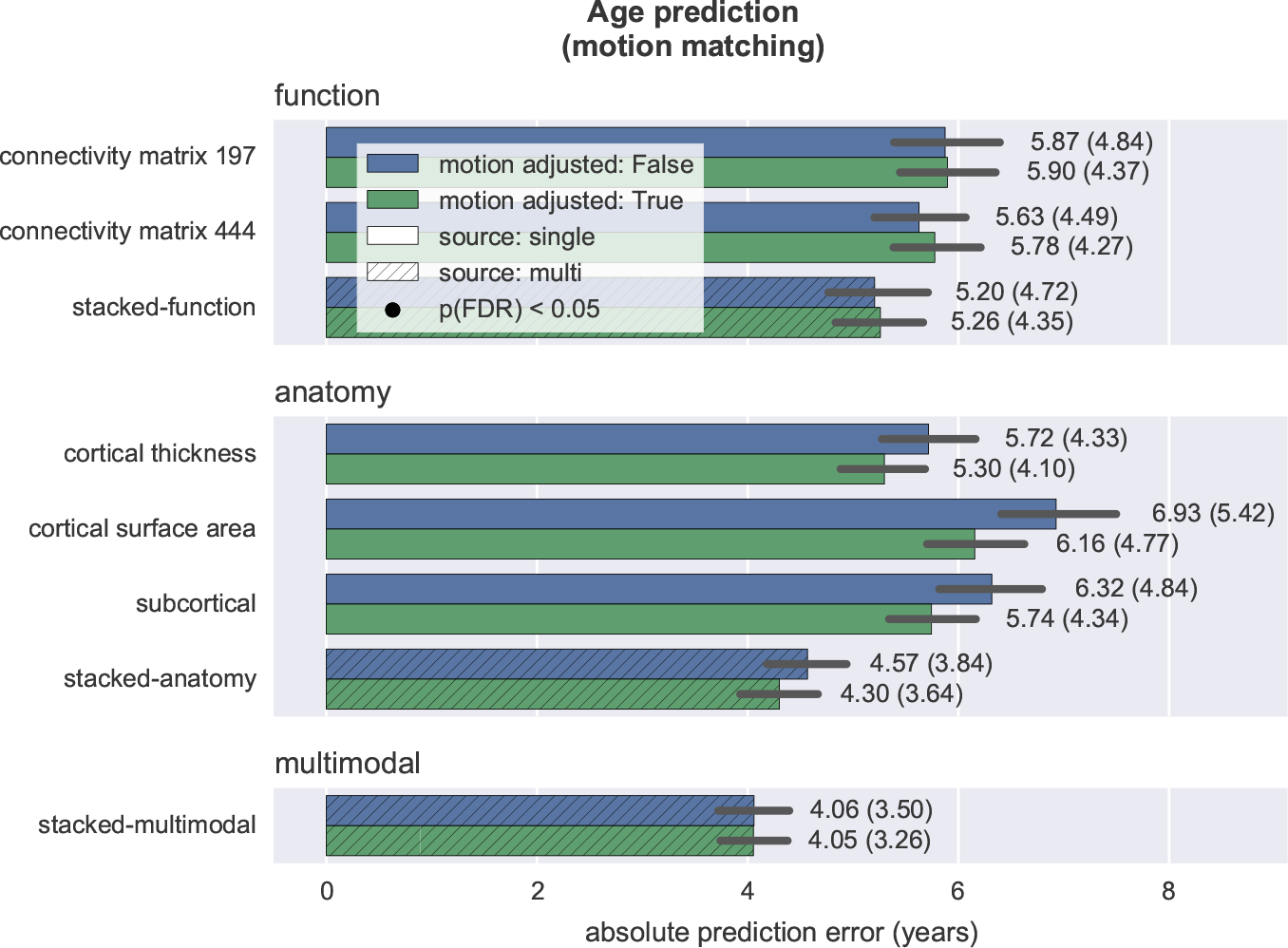
Motion adjustment via motion matching. Motion adjusted: True: sample without *age* × *motion* correlation; False: sample with preserved *age* × *motion* correlation. Note that for all models prediction with and without motion matching is equally good, indicating that the models’ predictions are not driven by head motion. For full statistics see Table B.8.

**Table B.8:** Motion adjustment via motion matching. Note that for all models prediction with and without motion matching is equally good, indicating that the models’ predictions are not driven by head motion. Motion adjusted: True: sample without *age* × *motion* correlation; False: sample with preserved *age* × *motion* correlation. (Mann-Whitney U test: motion adjusted False against True; N(False) = 387, N(True) = 387). See Figure B.9.

|  | motion adjusted False |  |  | motion adjusted True |  |  | U | p(FDR) |
| --- | --- | --- | --- | --- | --- | --- | --- | --- |
|  | M(APE) | SD(APE) | R2 | M(APE) | SD(APE) | R2 |  |  |
| connectivity matrix 197 | 5.87 | 4.83 | 0.70 | 5.90 | 4.36 | 0.61 | 7.3e+04 | 0.65 |
| connectivity matrix 444 | 5.63 | 4.48 | 0.73 | 5.78 | 4.26 | 0.63 | 7.3e+04 | 0.65 |
| stacked-function | 5.20 | 4.71 | 0.74 | 5.26 | 4.35 | 0.66 | 7.3e+04 | 0.65 |
| cortical thickness | 5.72 | 4.32 | 0.73 | 5.30 | 4.10 | 0.67 | 7.9e+04 | 0.49 |
| cortical surface area | 6.93 | 5.41 | 0.60 | 6.16 | 4.76 | 0.56 | 8e+04 | 0.49 |
| subcortical | 6.32 | 4.83 | 0.67 | 5.74 | 4.34 | 0.62 | 7.9e+04 | 0.49 |
| stacked-anatomy | 4.57 | 3.83 | 0.82 | 4.30 | 3.63 | 0.77 | 7.8e+04 | 0.65 |
| stacked-multimodal | 4.06 | 3.50 | 0.85 | 4.05 | 3.26 | 0.80 | 7.4e+04 | 0.86 |

### Appendix B.6. Generalization to new site

#### Appendix B.6.1. Generalization to new site: One sample training

**Figure B.10:**
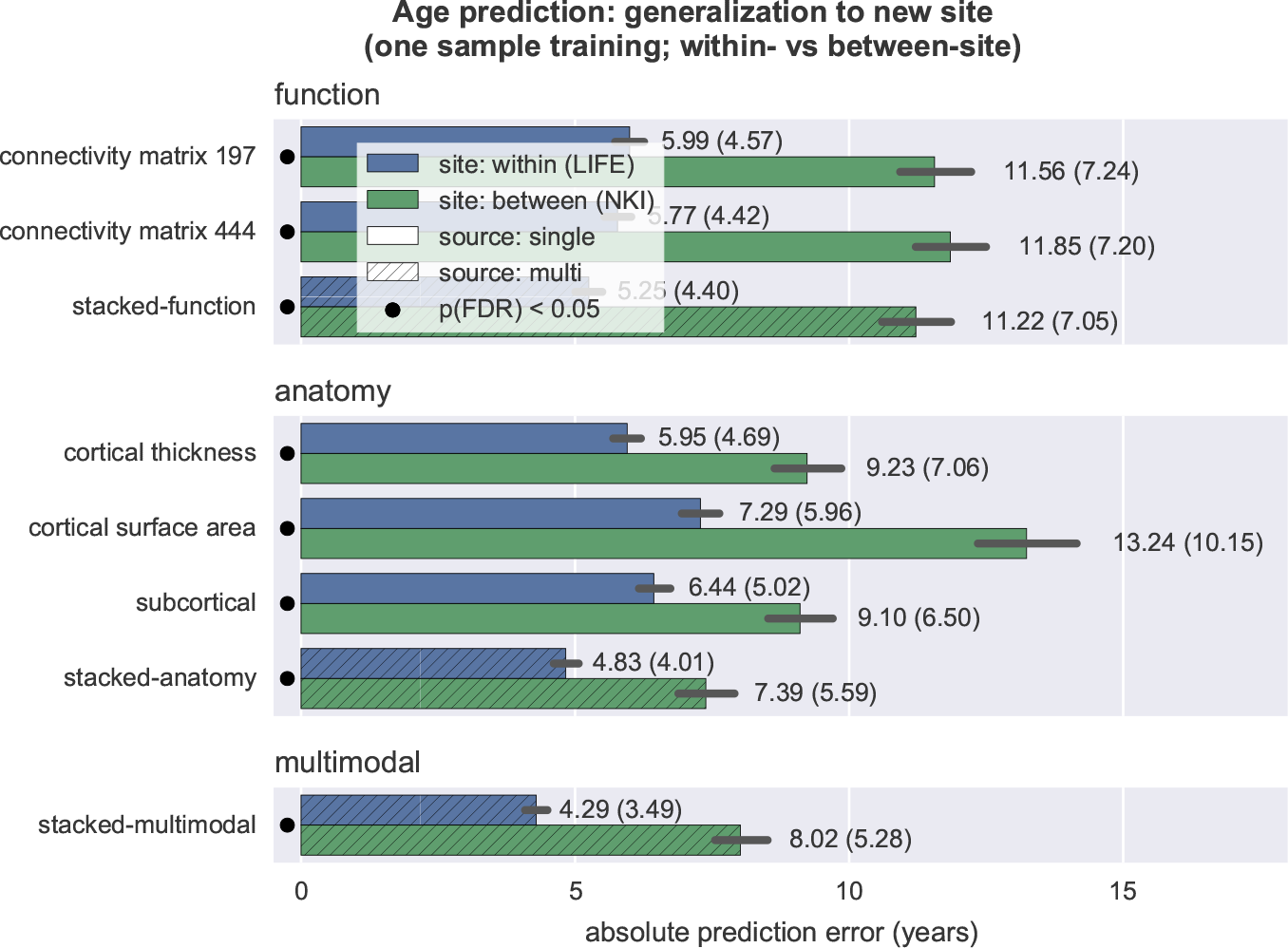
Standard training procedure (one sample training) showed significantly (*p*(*FDR*) < 0.05 as indicated by the black dot) better prediction performance in LIFE data (within site) than in NKI data (between site). See Table B.9).

**Table B.9:** Generalization to new site. Standard training procedure (one sample training) showed significantly better prediction performance in LIFE data (within site) than in NKI data (between site) (Mann-Whitney U test: site within (LIFE) against between (NKI); N(within (LIFE)) = 1177, N(between (NKI)) = 475). See Figure B.10.

|  | site within (LIFE) |  |  | site between (NKI) |  |  | U | p(FDR) |
| --- | --- | --- | --- | --- | --- | --- | --- | --- |
|  | M(APE) | SD(APE) | R2 | M(APE) | SD(APE) | R2 |  |  |
| connectivity matrix 197 | 5.99 | 4.57 | 0.75 | 11.56 | 7.23 | 0.48 | 1.5e+05 | 7.4e-49 |
| connectivity matrix 444 | 5.77 | 4.42 | 0.77 | 11.85 | 7.19 | 0.46 | 1.4e+05 | 1.7e-57 |
| stacked-function | 5.25 | 4.40 | 0.80 | 11.22 | 7.04 | 0.51 | 1.3e+05 | 5.4e-63 |
| cortical thickness | 5.95 | 4.69 | 0.75 | 9.23 | 7.06 | 0.62 | 2.1e+05 | 4.4e-17 |
| cortical surface area | 7.29 | 5.95 | 0.62 | 13.24 | 10.14 | 0.22 | 1.9e+05 | 8.3e-26 |
| subcortical | 6.44 | 5.02 | 0.71 | 9.10 | 6.49 | 0.65 | 2.1e+05 | 1.4e-14 |
| stacked-anatomy | 4.83 | 4.01 | 0.83 | 7.39 | 5.59 | 0.76 | 2e+05 | 7.5e-19 |
| stacked-multimodal | 4.29 | 3.49 | 0.87 | 8.02 | 5.27 | 0.74 | 1.6e+05 | 2.7e-43 |

#### Appendix B.6.2. Generalization to new site: Two samples training

**Table B.10:** Generalization to new site. Comparing test prediction performance on NKI data (between site); training on one vs two sites (Wilcoxon signed-rank test: training one sample against two samples; Ν = 429).

|  | training one sample |  |  | training two samples |  |  | W | p(FDR) |
| --- | --- | --- | --- | --- | --- | --- | --- | --- |
|  | M(APE) | SD(APE) | R2 | M(APE) | SD(APE) | R2 |  |  |
| connectivity matrix 197 | 11.53 | 7.30 | 0.48 | 10.35 | 7.15 | 0.56 | 3.3e+04 | 1.3e-06 |
| connectivity matrix 444 | 11.74 | 7.22 | 0.47 | 10.17 | 6.94 | 0.57 | 2.2e+04 | 9e-20 |
| stacked-function | 11.12 | 7.05 | 0.51 | 9.90 | 6.88 | 0.59 | 3.8e+04 | 0.0015 |
| cortical thickness | 9.21 | 7.03 | 0.62 | 8.11 | 5.93 | 0.72 | 3.3e+04 | 1.3e-06 |
| cortical surface area | 13.23 | 10.20 | 0.22 | 11.64 | 8.29 | 0.43 | 3.2e+04 | 3.5e-07 |
| subcortical | 9.04 | 6.50 | 0.65 | 8.01 | 6.18 | 0.71 | 3.8e+04 | 0.0012 |
| stacked-anatomy | 7.37 | 5.50 | 0.76 | 6.88 | 5.25 | 0.79 | 4.3e+04 | 0.17 |
| stacked-multimodal | 8.00 | 5.21 | 0.74 | 6.93 | 5.08 | 0.79 | 3.6e+04 | 0.00012 |

**Table B.11:** Generalization to new site. After training the model on a mixed-site sample (two sample training, *N*_*training,LIFE*_ = 1177; *N*_*training,NKI*_ = 46), predictions on the NKI data improve, but the predictions on the main training site LIFE (within site) still are significantly better than on the minor training site NKI (between site). (Mann-Whitney U test: site within (LIFE) against between (NKI); N(within (LIFE)) = 1177, N(between (NKI)) = 429). See Figure B.11.

|  | site within (LIFE) |  |  | site between (NKI) |  |  | U | p(FDR) |
| --- | --- | --- | --- | --- | --- | --- | --- | --- |
|  | M(APE) | SD(APE) | R2 | M(APE) | SD(APE) | R2 |  |  |
| connectivity matrix 197 | 6.51 | 5.03 | 0.71 | 10.35 | 7.15 | 0.56 | 1.7e+05 | 1.5e-21 |
| connectivity matrix 444 | 5.87 | 4.53 | 0.76 | 10.17 | 6.94 | 0.57 | 1.6e+05 | 5.8e-29 |
| stacked-function | 5.32 | 4.49 | 0.79 | 9.90 | 6.88 | 0.59 | 1.5e+05 | 1.7e-37 |
| cortical thickness | 6.27 | 4.88 | 0.73 | 8.11 | 5.93 | 0.72 | 2.1e+05 | 7.3e-08 |
| cortical surface area | 7.81 | 6.13 | 0.57 | 11.64 | 8.29 | 0.43 | 1.8e+05 | 1e-16 |
| subcortical | 7.91 | 6.50 | 0.55 | 8.01 | 6.18 | 0.71 | 2.5e+05 | 0.51 |
| stacked-anatomy | 5.11 | 4.27 | 0.81 | 6.88 | 5.25 | 0.79 | 2e+05 | 2.2e-09 |
| stacked-multimodal | 4.46 | 3.70 | 0.85 | 6.93 | 5.08 | 0.79 | 1.8e+05 | 1.8e-19 |

**Figure B.11:**
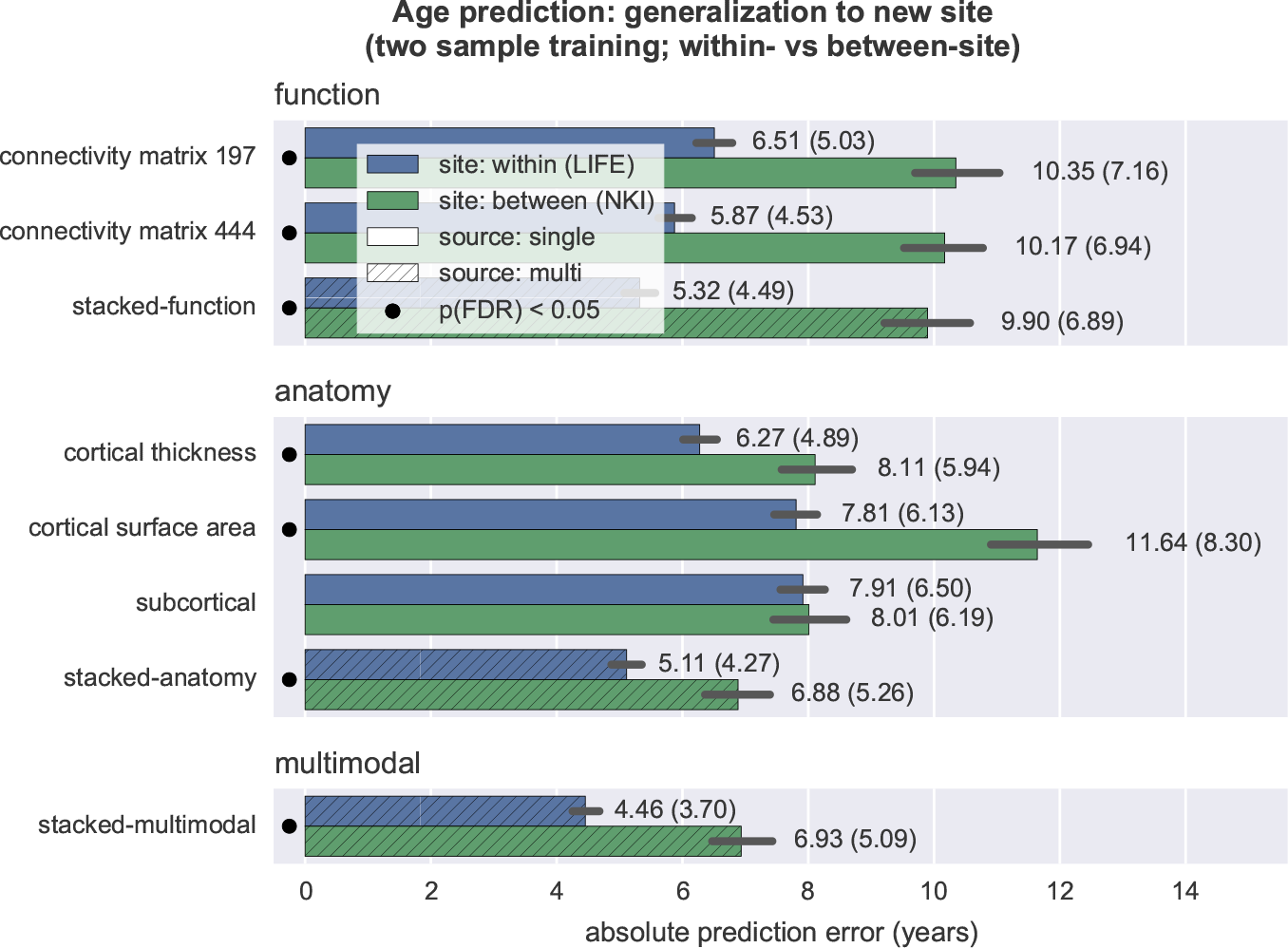
After training the model on a mixed-site sample (two sample training, *N*_*training,LIFE*_ = 1177; *N*_*training,NKI*_ = 46), predictions on the NKI data improve (Table B.10), but the predictions on the main training site LIFE (within site) still are significantly better than on the minor training site NKI (between site).

#### Appendix B.6.3. Generalization to new site: Training on full LIFE sample

**Table B.12:** Generalization to new site; training on full LIFE sample (*N*_*training,LIFE*_ = 2377). Comparing test prediction performance on NKI data (between site); training on one sample (LIFE sample only) vs two samples (LIFE + NKI samples; *Ν*_*training,ΝKI*_ = 46. (Wilcoxon signed-rank test: training one sample against two samples; Ν = 429). See Figure B.12.

|  | training one sample |  |  | training two samples |  |  | W | p(FDR) |
| --- | --- | --- | --- | --- | --- | --- | --- | --- |
|  | M(APE) | SD(APE) | R2 | M(APE) | SD(APE) | R2 |  |  |
| connectivity matrix 197 | 11.31 | 7.12 | 0.50 | 9.75 | 6.74 | 0.60 | 2.9e+04 | 1.2e-10 |
| connectivity matrix 444 | 11.05 | 7.17 | 0.51 | 9.45 | 6.50 | 0.63 | 2.5e+04 | 3.7e-16 |
| stacked-function | 10.27 | 6.59 | 0.58 | 8.88 | 6.41 | 0.66 | 3.7e+04 | 0.00028 |
| cortical thickness | 8.59 | 6.51 | 0.67 | 7.79 | 5.93 | 0.73 | 3.3e+04 | 4.2e-07 |
| cortical surface area | 12.08 | 9.24 | 0.35 | 11.08 | 7.89 | 0.48 | 3.5e+04 | 5.4e-05 |
| subcortical | 8.97 | 6.50 | 0.65 | 8.11 | 6.23 | 0.71 | 3.9e+04 | 0.0046 |
| stacked-anatomy | 7.35 | 5.29 | 0.77 | 6.74 | 4.94 | 0.80 | 4.2e+04 | 0.1 |
| stacked-multimodal | 7.79 | 4.96 | 0.76 | 6.56 | 4.49 | 0.82 | 3.5e+04 | 5.4e-05 |

**Figure B.12:**
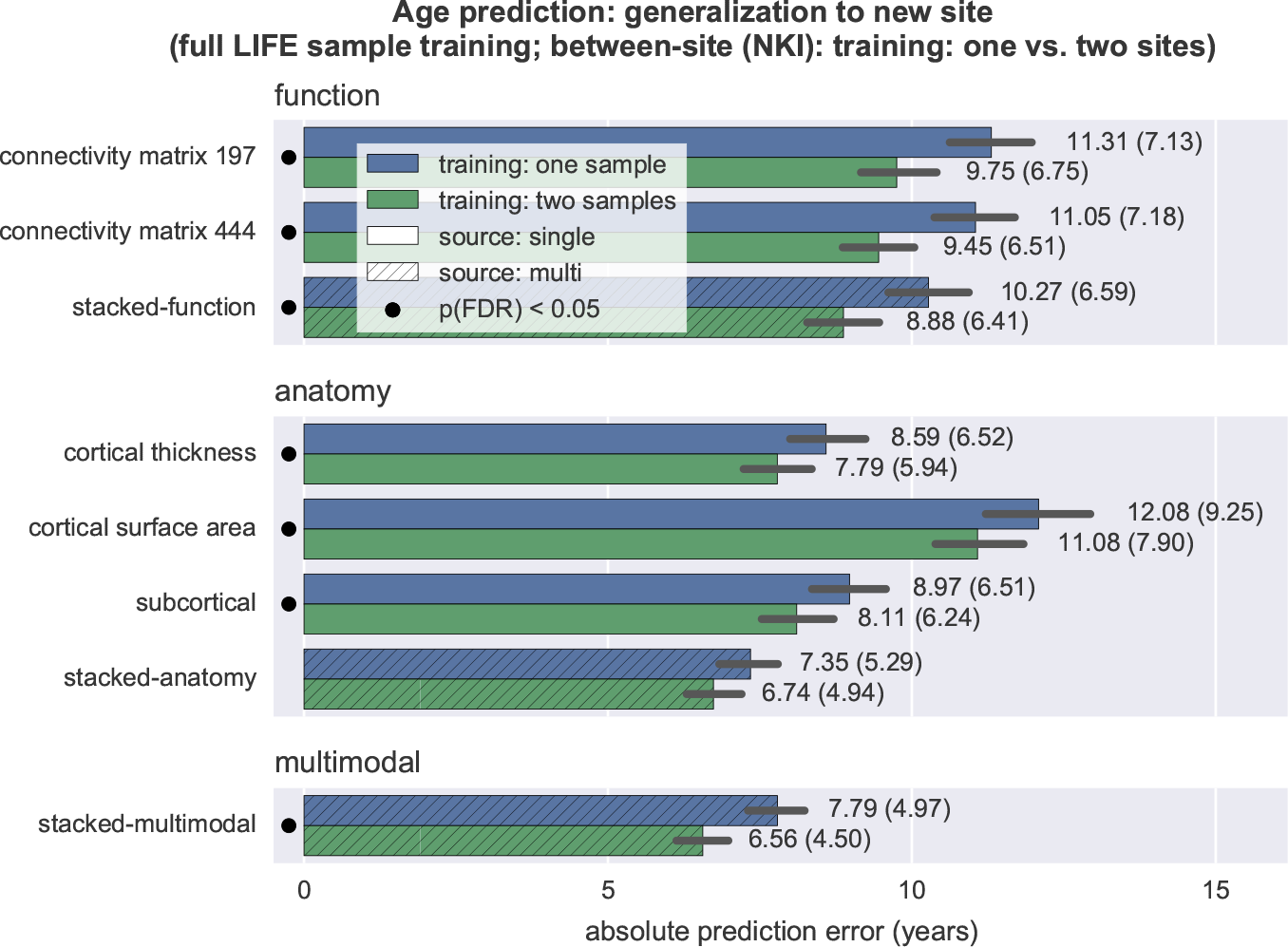
Generalization to new site; training on full LIFE sample *N*_*training,LIFE*_ = 2377). Comparing test prediction performance on NKI data (between site); training on one sample (LIFE sample only) vs two samples (LIFE + NKI samples; *N*_*training,NKI*_ = 46. See Table B.12.

**Figure B.13:**
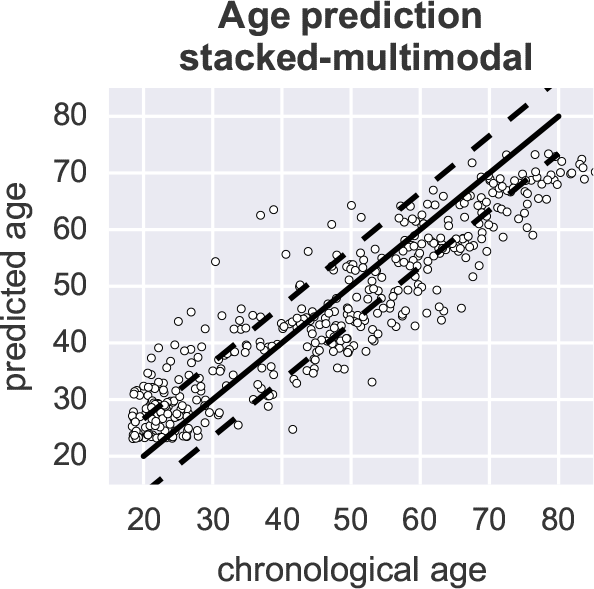
Chronological and predicted age for the NKI test set from the *stacked-multimodal* model with the two sample training approach. Circles represent subjects, the solid line the perfect prediction, dashed lines the mean absolute prediction error (6.56 years).

### Appendix B.7. Robustness of two sample training approach

**Table B.13:**
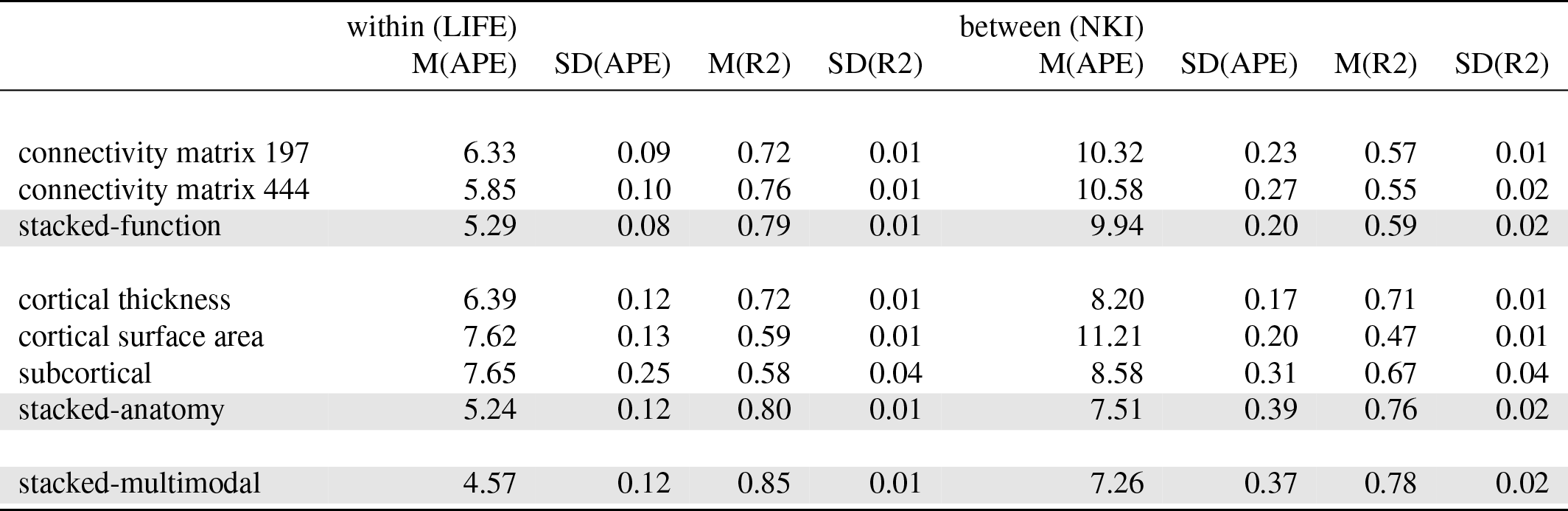
Stability of two sample training (cf. Table B.11) over ten random splits.

|  | within (LIFE) |  |  |  | between (NKI) |  |  |  |
| --- | --- | --- | --- | --- | --- | --- | --- | --- |
|  | M(APE) | SD(APE) | M(R2) | SD(R2) | M(APE) | SD(APE) | M(R2) | SD(R2) |
| connectivity matrix 197 | 6.33 | 0.09 | 0.72 | 0.01 | 10.32 | 0.23 | 0.57 | 0.01 |
| connectivity matrix 444 | 5.85 | 0.10 | 0.76 | 0.01 | 10.58 | 0.27 | 0.55 | 0.02 |
| stacked-function | 5.29 | 0.08 | 0.79 | 0.01 | 9.94 | 0.20 | 0.59 | 0.02 |
| cortical thickness | 6.39 | 0.12 | 0.72 | 0.01 | 8.20 | 0.17 | 0.71 | 0.01 |
| cortical surface area | 7.62 | 0.13 | 0.59 | 0.01 | 11.21 | 0.20 | 0.47 | 0.01 |
| subcortical | 7.65 | 0.25 | 0.58 | 0.04 | 8.58 | 0.31 | 0.67 | 0.04 |
| stacked-anatomy | 5.24 | 0.12 | 0.80 | 0.01 | 7.51 | 0.39 | 0.76 | 0.02 |
| stacked-multimodal | 4.57 | 0.12 | 0.85 | 0.01 | 7.26 | 0.37 | 0.78 | 0.02 |

